# Integrated Cell Landscape and Dynamics in the Progression of Bone Repair

**DOI:** 10.1101/2023.02.17.528986

**Authors:** Junfeng Shi, Jie Wang, Shi Yin, Sihan Lin, Fei Jiang, Maolin Zhang, Xiaolin Wu, Lijuan Shen, Xuefeng Gu, Ruomeng Yang, Jiawei Yang, Jin Wen, Wenjie Zhang, Qing Chang, Xinquan Jiang

## Abstract

Bone homeostasis and repair is a systematic progress with spatiotemporal interaction of multiple cell types involved in skeletal and immune system. Precise spatiotemporal regulation of cell type-specific functions in bone repair contributes to further development of tissue engineering and regenerative medicine. Here, we utilized single-cell RNA sequencing to illustrate a map of cell landscape and dynamics in the progression of rodent bone self-healing and a perturbation by lymphoid cell-deficiency. We identified different functions of myeloid cell and lymphoid cell to osteogenesis and angiogenesis during bone repair and their mutual complementation under lymphoid cell-deficient condition. Additionally, we used CD34^+^ humanized reconstituted mice to reveal further insights into the mechanism of human bone homeostasis and repair. Our integrated cellular analysis of bone repair explores the functional diversity and complementation between myeloid cells and lymphoid cells during bone healing process and provides further therapeutic implications for the treatment of bone disease and degeneration following ageing.

## Introduction

Tissue homeostasis and repair/regeneration originate from external cellular interplay and internal molecule interaction among different cell types (*1*). The dynamic and integrated cell network during tissue repair progression differs from time point and immune status (*2*). Precise spatiotemporal regulation of cell type-specific functions contributes to further development of tissue engineering and regenerative medicine (*3*). Bone homeostasis and repair have been keenly focused for its physical scaffold function and self-healing capacity (*4*). Once bone tissue suffered from a damage of micro-defects or fractures, repair and functional regeneration of this injury are primarily carried out by an autonomous osteogenic system (*5*), and can be accelerated by administration of medications or surgery (*6*).

Bone repair could also be impacted in the elderly, oncological and immunocompromised patients, mainly due to a disorder of the immune system (*4*). Previous studies demonstrated that bone repair process starts with an inflammatory reaction at the fracture site (*7*), whereas fracture healing would be accelerated at the early stage in the absence of the adaptive immune system (*8*), but delayed at the bone remodeling stage with deficiency of innate and adaptive immunity (*9*), suggesting a complex relationship between skeletal and immune system.

In recent years, cell therapy has gradually been highlighted for the non-surgical treatment of bone defect repair and fracture healing, and the research in related fields is also increasing. Potential regulating target of cells for bone injury repair mainly includes progenitor or stem cells for bone reformation (such as skeletal stem cells and mesenchymal stem cells) (*10*); (*11*)], stromal cells (such as fibroblasts and endothelial cells) (*12*) and immune cells that regulate bone healing and remodeling (*13*). It comes to a question that when and how those types of cells participate in bone healing process and whether they are available to the therapeutic application in bone disease.

With the help of single-cell sequencing technology, we can couple pathophysiological conditions and single-cell performance to enhance our understanding of cell dynamics in bone repair progression (*4*). In this study, we used the single-cell sequencing method to elucidate cellular activity and fate in the timeline of bone repair. At the same time, we used lymphoid cell-deficient and humanized reconstituted mice to explore the functional diversity and complementation between myeloid cells and lymphoid cells during bone healing process, which enables us to reveal key insights into the mechanism of bone homeostasis and repair, and provides further therapeutic implications for the treatment of bone disease and degeneration following ageing.

## Results

### A transcriptome atlas of bone repair at a single-cell level

In order to study the changes of cell types in the injury site during bone repair in mice, we first introduced a bone hole drilling model in the calvaria cortical region of ICR mice, which can build a stable reproducible environment without mechanical loading from animal gait, and make it possible to ascertain how cells contribute to the repair process by modelling facture healing and self-healing bone defect (drilled hole < 3mm in mice, n=18, **(*14*)**). According to the previous study on the temporal processes of bone healing in fractures and distraction osteogenesis of mice **(*15*); (*14*)**, we decided to harvest the tissues in the bone hole drilling site at day 7 (inflammatory response-callus formation), day 14 (endochondral formation), and day 21(bone remodeling) following the surgery (n = 6 at each time point, 3 were subjected to droplet-based scRNA-seq [Figure 1A], and 3 were subjected to micro-CT to observe the progression of bone repair [Figure 1B]). Hematoxylin/Eosin (HE) staining and fluorescence of calcein and alizarin red confirmed the bone reformation at 7d, 14d and 21d, respectively. The single-cell sequencing captured 9755, 10549 and 7875 cells on the 7d, 14d, and 21d of bone repair, which could be clustered into 21, 17, and 19 clusters, respectively, with the t-distributed Stochastic Neighborhood Embedding (t-SNE) method (Figure 1C).

**Figure 1.**
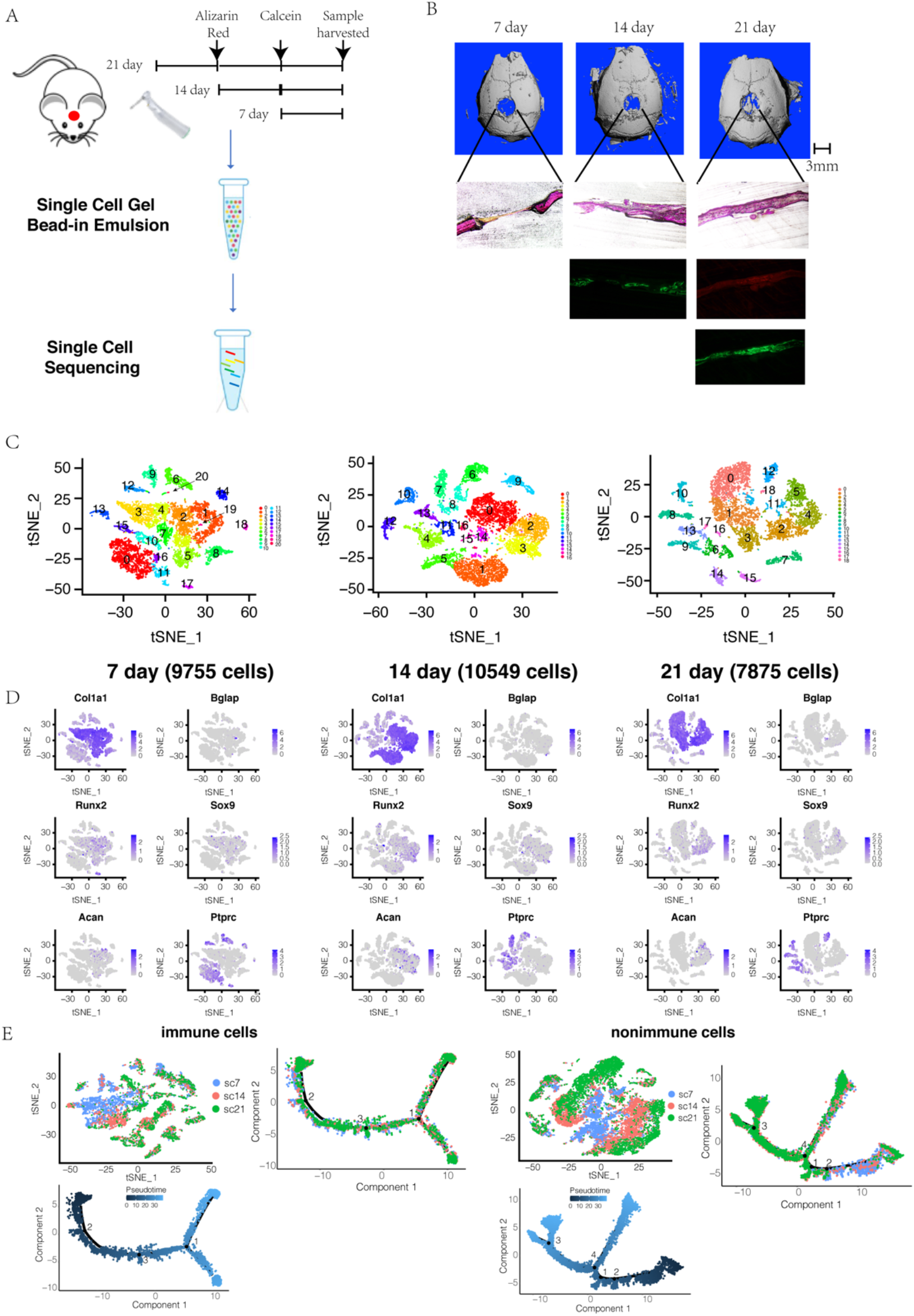
Cellular landscape and dynamics of bone repair progression under normal condition. (A) Study design. (B) Micro-CT image, Hematoxylin and eosin (H…E) staining, alizarin red S (red) and calcein (green) labeling of new bone formation and mineralization. (C) Cell census of cells in bone reformation site at 7 day (left), 14 day (middle), and 21 day (right) colored by cell subset from t-Stochastic Neighborhood Embedding (t-SNE). (D) Distribution of osteogenesis, chondrogenesis and immune marker genes in C (E) Cell census of immune cells (left) and non-immune cells in bone reformation site colored by post-surgery time points from t-SNE and pseudotime analysis.

Diffusion map analysis (DMA) was also performed **(*16*)**(Figures S1A-B). We confirmed the expression of osteogenesis/cartilage-related genes between different groups, and found that at 7d, cells expressing Col1a1 were widely distributed in cluster 1∼5, 7 and 8, while osteogenesis-related Runx2-expressed cells were mainly in clusters 1,2 and 7. Immune cells (Cd45, also called Ptprc^+^) were mainly in cluster 0, 10∼12, 14, 16, 17 (Figure 1C Left and 1D Left). Cells expressing cartilage-related genes (Sox9, Acan) have not yet appeared in large numbers. At day 14, Col1a1 and Runx2 were abundantly expressed in cells of cluster 0∼3, 5, 14, 16, and Cd45^+^ cells were mainly in cluster 4, 6∼8, 11, 13, and cells expressing Sox9 and Acan also appeared in cluster 0∼3 (Figure 1C Middle and 1D Middle). At day 21, cells expressing Col1a1 appeared in clusters 0∼5, while cells expressing Runx2 were mainly distributed in clusters 2 and 4. Cells expressing Sox9 and Acan were mainly in cluster 5, while the Cd45^+^ cells mainly appeared in clusters 6, 8, 9, 12, and 13 (Figure 1C Right and 1D Right).

In order to better study the types and functions of cells in different phases of bone repair, we fitted the cells at day 7, 14 and 21 together for analysis, and divided the cells into immune cells and non-immune cells according to the expression of Cd45 to perform independent studies (Figure 1E). By clustering with the t-SNE method and single-cell trajectory based on time point and pseudotime analysis (PSA) **(*17*)**, we confirmed that both immune cells and non-immune cells have independent distribution at day 7, 14 and 21(Figure 1E), and could be classified into 19 clusters, respectively (Figures S1C and S1E). By overlapping cluster-level relatedness based on cluster graph abstraction analysis (CGAA) **(*18*)** and the linkage distances based on the correlation coefficient between results of functional enrichment analysis in each cluster **(*19*)**, it was found that in the immune cell population, cluster 0-1-17 (cluster 0 : 7d&14d, microglial cells; cluster 1: 7d, macrophages; cluster 17: 7d&14d, dendritic cells (DCs), functions: phagocytosis, interleukin-6 production), cluster 5-6 (half of cells in cluster 5 in 7d, cells in cluster 6 were equidistributed in 7&14&21d, neutrophils, functions: neutrophil migration/chemotaxis/activation, apoptosis, exocytosis, interleukin-6 production), cluster 8-13 (half of cells in 14d, cluster 8: Treg cells; cluster 13: γδ T cells, functions: Treg/γδ T cell differentiation, T cell activation, immune response-cell surface receptor signaling) and cluster 2-14 (half of cluster-2 cells in 14d, cells in cluster 14 were equidistributed in 7&14&21d, B cells; functions: B cell proliferation/activation, immunoglobulin [Ig] production, immune response-cell surface receptor signaling, antigen processing and presentation) are relatively close in gene expression profile and Gene Ontology (GO)-based functional illustration (Table 1A, Figure S1D and Extended in supplementary for Fig S1D). Other clusters were DC (cluster 3, equidistribution, antigen processing and presentation), osteoclast (cluster 7, 7d, bone resorption/remodeling), Cd45^+^ fibroblast-like multipotential progenitors (cluster 9, 7/14d, p38-MAPK cascade, ossification, angiogenesis, BMP signaling pathway, Wnt signaling pathway, neurogenesis), NK cells (cluster 12, half in 14d, response to external biotic stimulus, NK cells-mediated immunity/cytotoxicity), DC (cluster 18, 7d, antigen processing and presentation).

**Table 1.** Cell clustering in Fig1/FigS1

**immune cell clustering**
| Cluster | Cell Type | Biomarker | Reference |
| --- | --- | --- | --- |
| Cluster 0 | microglial cell | Cx3cr1<sup>+</sup> |  |
| Cluster 1 | M2 macrophage | Mrc1<sup>+</sup>, Arg1<sup>+</sup> |  |
| Cluster 2 | B cell | Cd79a<sup>+</sup>, Cd79b<sup>+</sup> |  |
| Cluster 3 | dendritic cell | Cd209a<sup>+</sup>, Cd74<sup>+</sup> |  |
| Cluster 4 | B cell | Cd79a<sup>+</sup>, Vpreb1<sup>+</sup>, Vpreb3<sup>+</sup> |  |
| Cluster 5 | neutrophil | Csf3r<sup>+</sup> |  |
| Cluster 6 | neutrophil | Ly6g<sup>+</sup> |  |
| Cluster 7 | osteoclast | Ctsk<sup>+</sup> |  |
| Cluster 8 | Treg cell | Cd3g<sup>+</sup>, Cd4<sup>+</sup>, Foxp3<sup>+</sup> |  |
| Cluster 9 | fibroblast | Col5a2<sup>+</sup> |  |
| Cluster 10 | monocyte, proliferating | Ly6c2<sup>+</sup>, Mki67<sup>+</sup> |  |
| Cluster 11 | monocyte | Ly6c2<sup>+</sup> |  |
| Cluster 12 | NK cell | Ncr1<sup>+</sup> |  |
| Cluster 13 | gamma delta T cell | Trdc<sup>+</sup>, Cd3g<sup>+</sup> |  |
| Cluster 14 | B cell | Cd79a<sup>+</sup>, Cd79b<sup>+</sup> |  |
| Cluster 15 | plasmacytoid dendritic cell | Siglech<sup>+</sup>, Bst2<sup>+</sup> |  |
| Cluster 16 | stem cell | Ctla2a<sup>+</sup>, Kit<sup>+</sup> |  |
| Cluster 17 | dendritic cell | Cd11c<sup>+</sup> |  |
| Cluster 18 | dendritic cell | Cd74<sup>+</sup> |  |

**non-immune cell clustering**
| Cluster | Cell Type | Biomarker | Reference |
| --- | --- | --- | --- |
| Cluster 0 | fibroblast | Col5a2<sup>+</sup> |  |
| Cluster 1 | fibroblast | Col5a2<sup>+</sup> |  |
| Cluster 2 | fibroblast | Col5a2<sup>+</sup> |  |
| Cluster 3 | endothelial cell | Ly6c1<sup>+</sup>, Pi16<sup>+</sup> |  |
| Cluster 4 | pericyte | Acta2<sup>+</sup> |  |
| Cluster 5 | fibroblast | Col5a2<sup>+</sup> |  |
| Cluster 6 | endothelial cell | Ly6c1<sup>+</sup>, Pi16<sup>+</sup> |  |
| Cluster 7 | endothelial cell | Cdh5<sup>+</sup>, Pecam1<sup>+</sup> |  |
| Cluster 8 | smooth muscle cell | Rgs5<sup>+</sup> |  |
| Cluster 9 | fibroblast | Col5a2<sup>+</sup> |  |
| Cluster 10 | fibroblast | Col5a2<sup>+</sup> |  |
| Cluster 11 | erythrocyte | Car2<sup>+</sup> |  |
| Cluster 12 | fibroblast | Col5a2<sup>+</sup> |  |
| Cluster 13 | myofibroblast | Acta<sup>+</sup>, Myh11<sup>+</sup> |  |
| Cluster 14 | osteogenic cell | Bglap<sup>+</sup>, Sp7<sup>+</sup> |  |
| Cluster 15 | oligodendrocyte | Mbp<sup>+</sup>, Ptgsd<sup>+</sup> |  |
| Cluster 16 | erythrocyte | Car2<sup>+</sup> |  |
| Cluster 17 | endothelial cell | Ly6c1<sup>+</sup>, Pi16<sup>+</sup> |  |
| Cluster 18 | mural cell | Vtn<sup>+</sup>, Ifitm1<sup>+</sup> |  |

In the non-immune cell population, clusters 8-13 (cluster 8: 7&14d, smooth muscle cells; cluster 13: 14&21d, myofibroblast, function: actin filament organization, cell migration, cell locomotion, blood vessel development, muscle tissue development,) and cluster 2-5-6 (cluster 2: 14d, fibroblasts; cluster 5: 7d, fibroblasts; cluster 6: 21d, endothelial cells [ECs], functions: cell-cell adhesion, cell migration) were closer in gene expression and cellular function (Table 1B, Figure S1F and Extended in supplementary for Fig S1F). Other clusters were Ly6c1^+^ ECs (cluster 3, 21d, cell-cell adhesion, cell migration, tube morphogenesis), Chd5^+^ ECs (cluster 7, equidistribution, extracellular matrix organization, EC proliferation/differentiation, epithelial cell differentiation, cell-cell adhesion, mesenchymal cell differentiation, tube development, angiogenesis), osteogenic cells (cluster 14, 7&14d, endochondral ossification, osteoblast differentiation), oligodendrocytes (cluster 15, half in 14d, axon development, neurogenesis), mural cells (cluster 18, 7&14d, tube development, cell adhesion, cell locomotion, chemotaxis, cell junction organization)

### A single-cell level overview of bone repair progression in the absence of adaptive immune system

Bone repair could be impacted in the elderly, oncological and immunocompromised patients, mainly due to a deficiency of the immune system (*4*). In order to investigate how immunodeficiency influences bone healing systematically, we introduced the same hole drilling model to NPSG immunodeficient mice (Figure S2A). Like normal ICR mice, we obtained tissues from hole-drilled sites on day 7, 14, and 21 post-surgery for single-cell sequencing.

For the analysis of bone repair tissue under immunodeficiency on day 7, we obtained 21 clusters and DMA illustrated a cellular trend of differentiation (4794 cells, Figure 2A). Similarly, we split the cells into immune and non-immune cell populations based on their Cd45 expression, and compared them (named by n7) with the 7-day data of ICR mice (named by sc7), respectively. In the immune cell side, we identified 18 clusters (Figure 2B). Double-checked by single-cell trajectory based on PSA and DMA, respectively, we found that the cell status in the n7 group was significantly different from the sc7 group (Figures 2C and S2B). After analyzing the cluster-level relatedness and functional correlation coefficient-based linkage distances (Figures 2D-F), it was found that clusters 1-7 (cluster 1 in n7, cluster 7 in sc7, Ly6g^+^ neutrophils, response to bacterium, neutrophil activation, response to oxidative stress, neutrophil chemotaxis, apoptosis), cluster 6-9 (mainly in n7, granulocyte progenitor cells, function: not significant) and cluster 5-12 (cluster 5 in n7, cluster 12 in sc7, Csfr3^+^ neutrophils, functions: response to external biotic stimulus, neutrophil activation, response to oxidative stress, neutrophil chemotaxis, apoptosis, regulation of ossification, cell adhesion) are functionally close (Table 2A, Figure 2G and Extended in supplementary for Fig 2G). Other clusters were M2 macrophage (cluster 0, sc7; cluster 2, n7; response to external biotic stimulus, cell migration/locomotion), osteoclasts (cluster 4, sc7, bone resorption/remodeling), DCs (cluster 8, sc7, antigen processing and presentation), B cells (cluster 10, sc7, B cell activation, adaptive immune response), Cxcr6^+^Cd8^+^T cells (cluster 11, sc7, T cell activation, adaptive immune response), plasmacytoid DCs (cluster 14, sc7,), osteoblast-like fibroblast (cluster 15, sc7, tube development, cell motility, osteoblast differentiation, ossification), and microglial cells (cluster 16, n7, cell adhesion, MAPK cascade, cell communication, chemotaxis, apoptosis, tube development, cell migration, angiogenesis, gliogenesis, neurogenesis).

**Figure 2.**
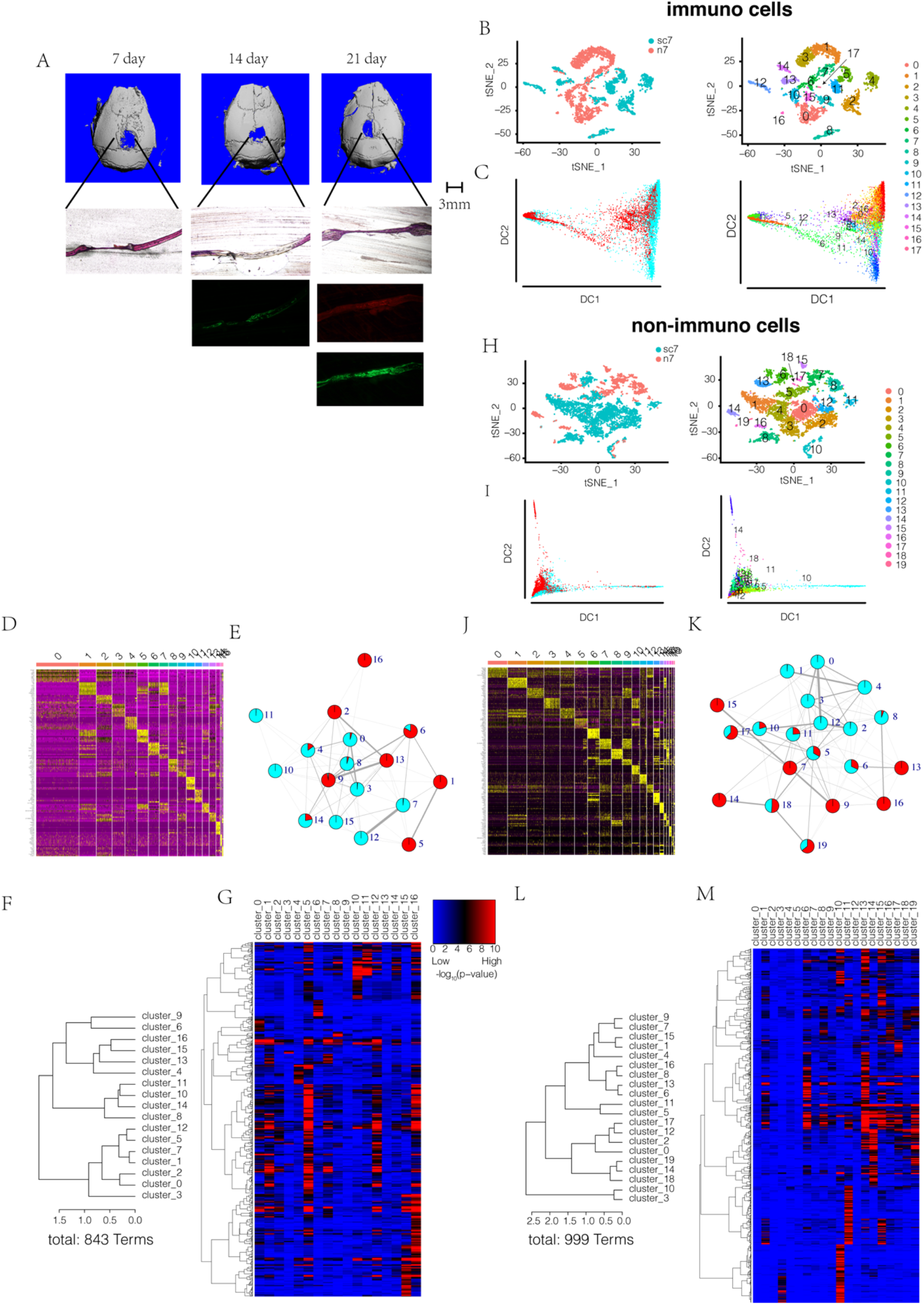
Cellular landscape and dynamics of bone repair progression under immune-deficient condition. (A) Micro-CT image, Hematoxylin and eosin (H…E) staining, alizarin red S (red) and calcein (green) labeling of new bone formation and mineralization. (B and C) Cell census of immune cells in bone reformation site colored by post-surgery time points and cell subset from t-SNE and diffusion map analysis. (D) Cluster signature genes of immune cells. Expression (row-wide Z score of ln(TP10K+1)) of top differentially expressed genes (rows) across the cells (columns) in each cluster. (E) Relatedness (edges, width indicates strength) between clusters (nodes, colored as post-surgery time points) based on cluster graph abstraction of B. (F) Cluster dendrogram of cell subsets in B based on the patterns of Gene Ontology (GO) enrichment; distance method: correlation; agglomeration method: Ward’s method. The linkage distance is the measure of similarity between cell subsets, with increasing distance representing decreasing degrees of similarity between samples. (G) GO heatmap and clustering dendrogram with GO terms along the horizontal axis. The heatmap was generated across all cell subsets. P values were calculated by the hypergeometric test and denoted by -log_10_P; overrepresented GO terms were shown in each cell subsets and colored by red and blue in Extend Figure-2G; distance method: correlation; agglomeration method: Ward’s method. (H and I) Cell census of non-immune cells in bone reformation site colored by post-surgery time points and cell subset from t-SNE and diffusion map analysis. (J) Cluster signature genes of non-immune cells. Expression (row-wide Z score of ln(TP10K+1)) of top differentially expressed genes (rows) across the cells (columns) in each cluster. (K) Relatedness (edges, width indicates strength) between clusters (nodes, colored as post-surgery time points) based on cluster graph abstraction of H. (L) Cluster dendrogram of cell subsets in H based on the patterns of GO enrichment; distance method: correlation; agglomeration method: Ward’s method. The linkage distance is the measure of similarity between cell subsets, with increasing distance representing decreasing degrees of similarity between samples. (M) GO heatmap and clustering dendrogram with GO terms along the horizontal axis. The heatmap was generated across all cell subsets. P values were calculated by the hypergeometric test and denoted by -log_10_P; overrepresented GO terms were shown in each cell subsets and colored by red and blue in Extend Figure-2M; distance method: correlation; agglomeration method: Ward’s method.

**Table 2.** Cell clustering in Fig2/FigS2

**immune cell clustering**
| Cluster | Cell Type | Biomarker | Reference |
| --- | --- | --- | --- |
| Cluster 0 | macrophage | Cd206<sup>+</sup>, Cd32<sup>+</sup> |  |
| Cluster 1 | neutrophils | Ly6g<sup>+</sup> |  |
| Cluster 2 | macrophage | Cd206<sup>+</sup>, Cd32<sup>+</sup> |  |
| Cluster 3 | monocytes | Ly6c2<sup>+</sup> |  |
| Cluster 4 | osteoclast | Ctsk<sup>+</sup> |  |
| Cluster 5 | stage II neutrophils | Csfr3<sup>+</sup> |  |
| Cluster 6 | stem cell | Prtn3<sup>+</sup>, Mpo<sup>+</sup> |  |
| Cluster 7 | neutrophils | Ly6g<sup>+</sup> |  |
| Cluster 8 | dendritic cell | Cd74<sup>+</sup> |  |
| Cluster 9 | stem cell | Prtn3<sup>+</sup>, Mpo<sup>+</sup> |  |
| Cluster 10 | B cell | Cd79a<sup>+</sup> |  |
| Cluster 11 | Cd8<sup>+</sup> T cell | Cd3g<sup>+</sup>, Cxcr6<sup>+</sup>, Cd8<sup>+</sup> |  |
| Cluster 12 | stage II neutrophils | Csfr3<sup>+</sup> |  |
| Cluster 13 | monocytes | Ly6c<sup>+</sup> |  |
| Cluster 14 | plasmacytoid dendritic cell | Siglech<sup>+</sup> |  |
| Cluster 15 | osteoblast-like fibroblast | Col5a2<sup>+</sup>, Bmp2k<sup>+</sup> |  |
| Cluster 16 | microglia cell | Cx3cr1<sup>+</sup> |  |

**non-immune cell clustering**
| Cluster | Cell Type | Biomarker | Reference |
| --- | --- | --- | --- |
| Cluster 0 | fibroblast | Col5a2<sup>+</sup> |  |
| Cluster 1 | endothelial cell | Ly6c1<sup>+</sup>, Pi16<sup>+</sup> |  |
| Cluster 2 | fibroblast | Col5a2<sup>+</sup> |  |
| Cluster 3 | pericyte cell, proliferating | Acta2<sup>+</sup>, Mki67<sup>+</sup> |  |
| Cluster 4 | fibroblast | Col5a2<sup>+</sup> |  |
| Cluster 5 | pericyte cell | Acta2<sup>+</sup> |  |
| Cluster 6 | endothelial cell | Cdh5<sup>+</sup>, Pecam1<sup>+</sup> |  |
| Cluster 7 | fibroblast | Col5a2<sup>+</sup>, Ecrg4<sup>+</sup>, Lgals3<sup>+</sup> |  |
| Cluster 8 | pericyte | Acta2<sup>+</sup> |  |
| Cluster 9 | fibroblast | Col5a2<sup>+</sup>, Ecrg4<sup>+</sup>, Lgals3<sup>+</sup> |  |
| Cluster 10 | erythroid cell | Alas2<sup>+</sup> |  |
| Cluster 11 | microglial cell | Cx3cr1<sup>+</sup> |  |
| Cluster 12 | chondrogenic cell | Comp<sup>+</sup>, Acan<sup>+</sup>, Col11a1<sup>+</sup> |  |
| Cluster 13 | endothelial cell | Cdh5<sup>+</sup>, Pecam1<sup>+</sup> |  |
| Cluster 14 | oligodendrocyte | Mbp<sup>+</sup>, Ptgsd<sup>+</sup> |  |
| Cluster 15 | fibroblast | Col5a2<sup>+</sup>, Ecrg4<sup>+</sup>, Lgals3<sup>+</sup> |  |
| Cluster 16 | pericyte | Acta2<sup>+</sup> |  |
| Cluster 17 | osteogenic cell | Runx2<sup>+</sup>, Bglap<sup>+</sup> |  |
| Cluster 18 | neuron-like epithelium cells | Cd44<sup>+</sup>, Enpp2<sup>+</sup> |  |
| Cluster 19 | oligodendrocyte | Mbp<sup>+</sup>, Mal<sup>+</sup> |  |

At the same time, we used weighted correlation network analysis (WGCNA) to investigate the hub genes contributing to differences of immune cells between sc7 and n7 (Figures S2C and S2D), and found that five modules were associated with phenotype and clustering, respectively (MEgreen and MEbrown was negatively correlated with phenotype, MEmagenta, MEblack and MEpink are positively correlated with clustering, Figure S2E). By extracting genes in each module, we found that genes in MEgreen were mainly involved in cluster 2, 6, 9, 13, 16; MEbrown in cluster 1, 5, 12, 13; MEmagenta in cluster 11; MEblack in cluster 14; MEpink in cluster 10, Figure S2F), indicating neutrophils in cluster 1, 5 (n7), granulocyte progenitor cells in cluster 6,9 (n7), macrophage in cluster 2 (n7), and microglial cells in cluster 16 (n7) play a role in immunodeficiency-induced bone repair disorder in the early stage.

In the non-immune cell group, we identified 20 clusters (Figure 2H). By using PSA and DMA-based single-cell trajectory, we found that n7 group was significantly different from sc7 group (Figures 2I and S2G). Then, we calculated the cluster-level relatedness (Figures 2J and 2K) and functional correlation coefficient-based linkage distances (Figure 2L), and found that cluster 7-9-15 (n7, fibroblast, Erk1/2 cascade, response to interleukin-1, tube development, cell migration/locomotion), cluster 14-18-19 (cluster 14: n7, oligodendrocyte; cluster 18: n7&sc7, neuron-like epithelium cells, cluster 19: n7&sc7, oligodendrocyte; neuron differentiation, neurogenesis, axonogenesis, oligodendrocyte differentiation, glial cell differentiantion), cluster 8-16 (cluster 8: sc7, cluster 16: n7, pericyte, smooth muscle cell proliferation, cell adhesion, tube development, angiogenesis, vasculogenesis), and cluster 6-13 (cluster 6: n7&sc7, cluster 13: n7, ECs, angiogenesis, epithelial cell differentiation/migration, cell adhesion, vasculature development, tight junction assembly, EC proliferation, cell migration) have similarity with each other (Table 2B, Figure 2M and Extended in supplementary for Fig 2M). Other clusters were ECs (cluster 1, sc7, cell migration/locomotion, tube morphogenesis), chondrogenic cells (cluster 12, sc7, chondrocyte differentiation), osteogenic cells (cluster 17, sc7&n7, osteoblast differentiation, ossification).

Likewise, we used WGCNA to look for the difference of non-immune cells between SC and N group (Figures S2H and S2I), and found that one module was associated with phenotype and clustering respectively (MEred was negatively correlated with phenotype and positively correlated with clustering) (Figure S2J). By extracting the genes in this module, we found that they are mainly in cluster 7, 9, 13, 14, 15, 16, 18, Figure S2K), indicating that Ecrg4^+^Lgals3^+^ fibroblast, Cdh5^+^ endothelial cell, Ptgds^+^ oligodendrocyte, Acta2^+^ pericyte and Enpp2^+^ neuron-like epithelium cell contributed to the bone repair at the early stage, and could be functionally changed by immunodeficiency.

### Functional difference of immune cells during bone repair at homeostasis and under adaptive immune system-deficient conditions

According to the analysis of bone repair site under immunodeficiency at day 14, we obtained 18 clusters and performed DMA on them (4324 cells, Figure 3A, named by n14). Similarly, we split the cells into immune cell populations and non-immune cell populations based on their Cd45 expression and compared with the 14-day data of ICR mice (sc14). In the immune cell group, we identified 20 clusters (Figure 3B). PSA and DMA confirmed the different status between n14 group and sc14 group (Figures 3C and S3A). After analyzing the similarity between clusters (Figure 3D-F), it was found that cluster 16-17 (cluster 16: n14&sc14, cluster 17: sc14, fibroblasts, angiogenesis, ossification, cell junction organization, neuron development, cell migration/motility), cluster 7-9 (n14, neutrophils), cluster 8-15 (both mainly in sc14) and cluster 14-19 (cluster 14 mainly in n14, cluster 19 mainly in sc14) are relatively close. The detailed cell identities and GO-based functions are shown in Table 3A, Figure 3G and Extended file for Fig 3G, respectively.

**Figure 3.**
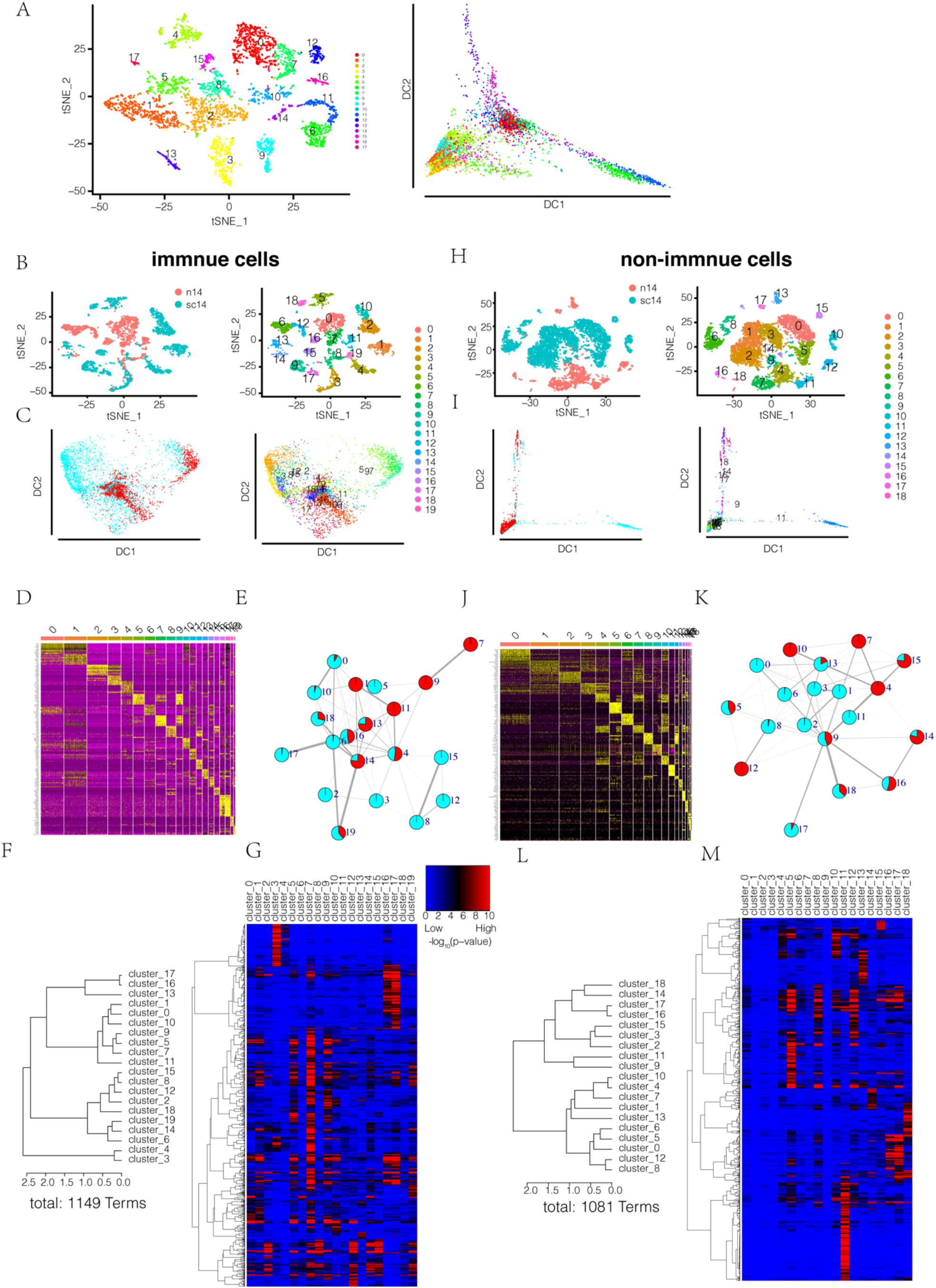
Immune cells present different functions during bone repair at homeostasis and under immuno-deficient conditions. (A) Cell census of post-surgery 14 day at bone reformation site colored by cell subsets from t-SNE and diffusion map analysis. (B and C) Cell census of immune cells in bone reformation site colored by post-surgery time points and cell subset from t-SNE and diffusion map analysis. (D) Cluster signature genes of immune cells. Expression (row-wide Z score of ln(TP10K+1)) of top differentially expressed genes (rows) across the cells (columns) in each cluster. (E) Relatedness (edges, width indicates strength) between clusters (nodes, colored as post-surgery time points) based on cluster graph abstraction of B. (F) Cluster dendrogram of cell subsets in B based on the patterns of Gene Ontology (GO) enrichment; distance method: correlation; agglomeration method: Ward’s method. The linkage distance is the measure of similarity between cell subsets, with increasing distance representing decreasing degrees of similarity between samples. (G) GO heatmap and clustering dendrogram with GO terms along the horizontal axis. The heatmap was generated across all cell subsets. P values were calculated by the hypergeometric test and denoted by -log_10_P; overrepresented GO terms were shown in each cell subsets and colored by red and blue in Extend Figure-3G; distance method: correlation; agglomeration method: Ward’s method. (H and I) Cell census of non-immune cells in bone reformation site colored by post-surgery time points and cell subset from t-SNE and diffusion map analysis. (J) Cluster signature genes of non-immune cells. Expression (row-wide Z score of ln(TP10K+1)) of top differentially expressed genes (rows) across the cells (columns) in each cluster. (K) Relatedness (edges, width indicates strength) between clusters (nodes, colored as post-surgery time points) based on cluster graph abstraction of H. (L) Cluster dendrogram of cell subsets in H based on the patterns of GO enrichment; distance method: correlation; agglomeration method: Ward’s method. The linkage distance is the measure of similarity between cell subsets, with increasing distance representing decreasing degrees of similarity between samples. (M) GO heatmap and clustering dendrogram with GO terms along the horizontal axis. The heatmap was generated across all cell subsets. P values were calculated by the hypergeometric test and denoted by -log_10_P; overrepresented GO terms were shown in each cell subsets and colored by red and blue in Extend Figure-3M; distance method: correlation; agglomeration method: Ward’s method.

**Table 3.** Cell clustering in Fig3/Fig3S

| A immune cell clustering |  |  |  |
| --- | --- | --- | --- |
| Cluster | Cell Type | Biomarker | Reference |
| Cluster 0 | macrophage | Adgre1<sup>+</sup> |  |
| Cluster 1 | macrophage | Adgre1<sup>+</sup> |  |
| Cluster 2 | B cell | Cd79a<sup>+</sup> |  |
| Cluster 3 | B cell | Cd79a<sup>+</sup> |  |
| Cluster 4 | neutrophil | Prtn3<sup>+</sup>, Mpo<sup>+</sup> |  |
| Cluster 5 | neutrophil | Ly6g<sup>+</sup>, Mmp9<sup>+</sup> |  |
| Cluster 6 | monocyte | Ccr2<sup>+</sup> |  |
| Cluster 7 | neutrophil | Csf3r<sup>+</sup> |  |
| Cluster 8 | T cell | Cd3g<sup>+</sup> |  |
| Cluster 9 | neutrophil | Ly6g<sup>+</sup> |  |
| Cluster 10 | macrophage | Adgre1<sup>+</sup> |  |
| Cluster 11 | monocytic MDSC | Ly6c<sup>+</sup>, Cd11b<sup>+</sup> |  |
| Cluster 12 | NK cell | Ncr1<sup>+</sup> |  |
| Cluster 13 | neutrophil | Prtn3<sup>+</sup>, Mpo<sup>+</sup> |  |
| Cluster 14 | dendritic cell | Cd74<sup>+</sup> |  |
| Cluster 15 | T cell | Cd3g<sup>+</sup> |  |
| Cluster 16 | fibroblast | Col5a2<sup>+</sup> |  |
| Cluster 17 | fibroblast | Col5a2<sup>+</sup> |  |
| Cluster 18 | plasmacytoid dendritic cell | Siglech<sup>+</sup> |  |
| Cluster 19 | dendritic cell | Cd74<sup>+</sup> |  |

| B non-immune cell clustering |  |  |  |
| --- | --- | --- | --- |
| Cluster | Cell Type | Biomarker | Reference |
| Cluster 0 | fibroblast | Col5a2<sup>+</sup> |  |
| Cluster 1 | fibroblast | Col5a2<sup>+</sup> |  |
| Cluster 2 | fibroblast | Col5a2<sup>+</sup> |  |
| Cluster 3 | fibroblast | Col5a2<sup>+</sup> |  |
| Cluster 4 | fibroblast | Col5a2<sup>+</sup>, Ecrg4<sup>+</sup> |  |
| Cluster 5 | BMEC | Cdh5<sup>+</sup> |  |
| Cluster 6 | fibroblast | Col5a2<sup>+</sup> |  |
| Cluster 7 | fibroblast | Col5a2<sup>+</sup>, Ecrg4<sup>+</sup> |  |
| Cluster 8 | pericyte | Acta2<sup>+</sup> |  |
| Cluster 9 | fibroblast | Col5a2<sup>+</sup> |  |
| Cluster 10 | fibroblast | Col5a2<sup>+</sup>, Ecrg4<sup>+</sup> |  |
| Cluster 11 | erythrocyte | Car2<sup>+</sup> |  |
| Cluster 12 | pericyte | Acta2<sup>+</sup> |  |
| Cluster 13 | stage I neutrophil | G0s2<sup>+</sup>, S100a8<sup>+</sup>, S100a9<sup>+</sup> |  |
| Cluster 14 | neuron-like epithelium cell | Cd44<sup>+</sup>, Enpp2<sup>+</sup> |  |
| Cluster 15 | osteogenic cell | Bglap<sup>+</sup> |  |
| Cluster 16 | oligodendrocyte | Mbp<sup>+</sup>, Ptgsd<sup>+</sup> |  |
| Cluster 17 | oligodendrocyte | Mbp<sup>+</sup>, Ptgsd<sup>+</sup> |  |
| Cluster 18 | neuroendocrine cell | Chgb<sup>+</sup> |  |

Next, By WGCNA (Figures S3B and S3C), we identified five modules associated with phenotype and clustering (MEblack and MEyellow were negatively correlated with phenotype, MEred and phenotype is positively correlated, MEturquoise and MEgrey are positively correlated with clustering, Figure S3D), respectively. Genes in each module were mainly distributed in cluster 1, 4, 7, 9, 11, 13, 14, 16 (MEblack); cluster 1, 7, 9, 11, 14 (MEyellow); cluster 2, 3 (MEred); cluster 16, 17 (MEturquoise); all cluster (MEgrey) (Figure S3E), indicating that during endochondral osteogenesis of bone repair, lymphoid cells, especially T cells and NK cells, started to be engaged in.

Towards the non-immune cell group, we identified 19 clusters (Figure 3H) and illustrated the difference of n14 from sc14 by PSA and DMA (Figures 3I and S3F). Cluster-level relatedness and functional correlation coefficient-based linkage distances identified the similarities of cluster 4-7-10 (mainly in n14), cluster 8-12 (cluster 8 mainly in sc14, cluster 12 mainly in n14) (Figure 3J-L). The detailed cell identities and functions are shown in Table 3B, Figure 3G and Extended file for Fig 3G. WGCNA showed that MEgreen was negatively correlated with phenotype and positively correlated with clustering (Figures S3G-I). Genes in MEgreen are mainly in cluster 4, 5, 7, 10, 12, 14, 15 (Figure S3J), indicating Ecrg4^+^fibroblasts, Cdh5^+^endothelial cells, Acta2^+^pericytes and Enpp2^+^epithelium cells were still working at an important site for endochondral osteogenesis of bone repair.

### Functional difference of non-immune cells during bone repair at homeostasis and under adaptive immune system-deficient conditions

For the analysis of bone repair site under immunodeficiency on day 21, we obtained 17 clusters and performed DMA on them (5358 cells, n21, Figure 4A). The cells were split into immune cell populations and non-immune cell populations by Cd45 expression, and compared with the 21-day data of ICR mice (sc21), respectively. In the immune cell population, we identified 20 clusters (Figure 4B) and showed the difference of the two groups using PSA and DMA, respectively (Figures 4C and S4A). After analyzing the similarity between clusters (Figure 4D-F), it was found that cluster 7-17 (cluster 7 mainly in n21, cluster 17 mainly in sc21), cluster 4-13-16-18 (cluster 4,13,18 mainly in sc21, cluster 16 in both groups), cluster 0-1-2 (mainly in n21), cluster 14-15 (cluster mainly in n21, cluster 15 mainly in sc21), cluster 5-6 (mainly in n21) were functionally close. The detailed cell identities and GO-based functions could be found in Table 4A, Figure 4G and Extended file for Fig 4G.

**Figure 4.**
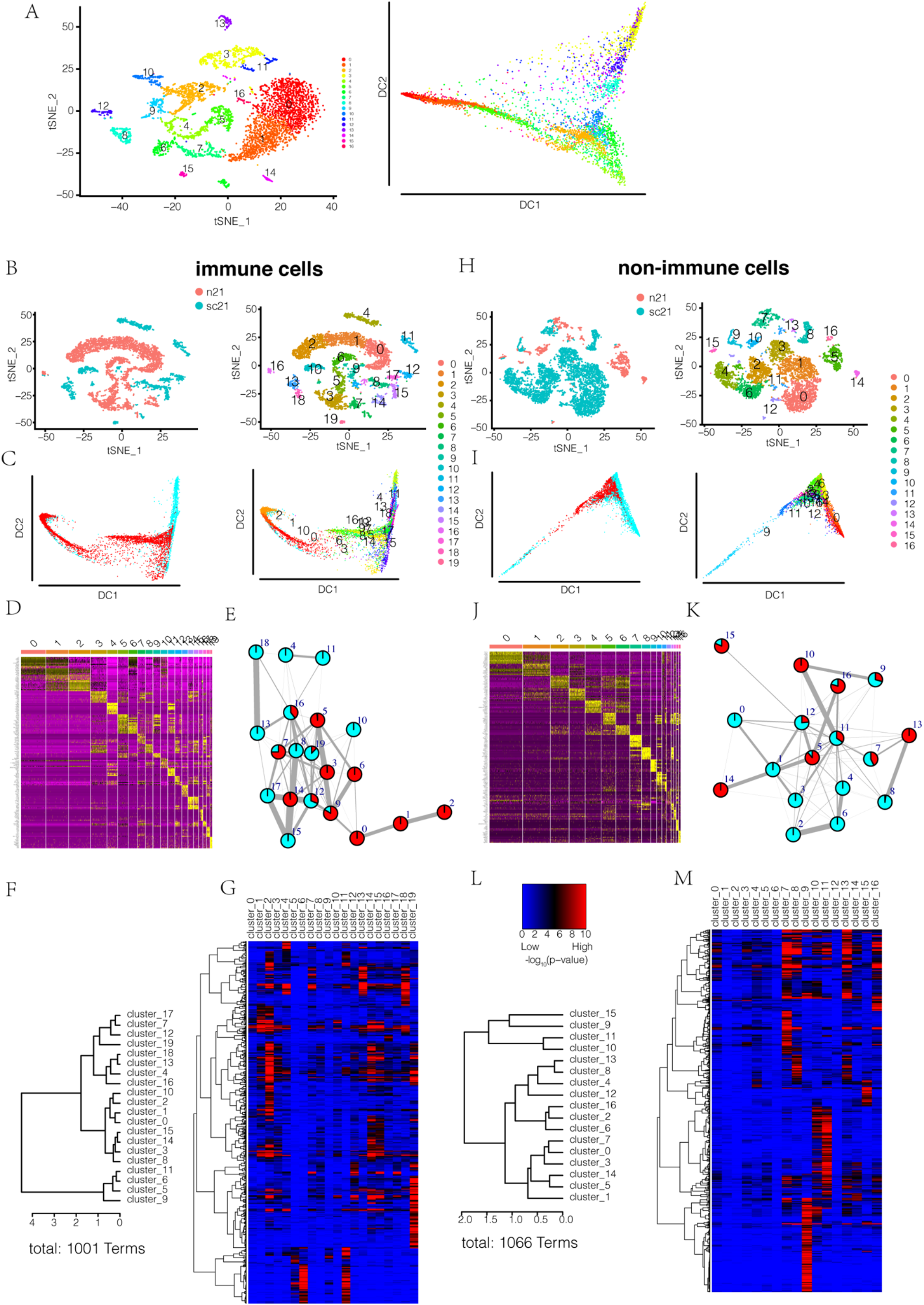
Non-immune cells present different functions during bone repair at homeostasis and under immuno-deficient conditions. (A) Cell census of post-surgery 14 day at bone reformation site colored by cell subsets from t-SNE and diffusion map analysis. (B and C) Cell census of immune cells in bone reformation site colored by post-surgery time points and cell subset from t-SNE and diffusion map analysis. (D) Cluster signature genes of immune cells. Expression (row-wide Z score of ln(TP10K+1)) of top differentially expressed genes (rows) across the cells (columns) in each cluster. (E) Relatedness (edges, width indicates strength) between clusters (nodes, colored as post-surgery time points) based on cluster graph abstraction of B. (F) Cluster dendrogram of cell subsets in B based on the patterns of GO enrichment; distance method: correlation; agglomeration method: Ward’s method. The linkage distance is the measure of similarity between cell subsets, with increasing distance representing decreasing degrees of similarity between samples. (G) GO heatmap and clustering dendrogram with GO terms along the horizontal axis. The heatmap was generated across all cell subsets. P values were calculated by the hypergeometric test and denoted by -log_10_P; overrepresented GO terms were shown in each cell subsets and colored by red and blue in Extend Figure-4G; distance method: correlation; agglomeration method: Ward’s method. (H and I) Cell census of non-immune cells in bone reformation site colored by post-surgery time points and cell subset from t-SNE and diffusion map analysis. (J) Cluster signature genes of non-immune cells. Expression (row-wide Z score of ln(TP10K+1)) of top differentially expressed genes (rows) across the cells (columns) in each cluster. (K) Relatedness (edges, width indicates strength) between clusters (nodes, colored as post-surgery time points) based on cluster graph abstraction of H. (L) Cluster dendrogram of cell subsets in H based on the patterns of GO enrichment; distance method: correlation; agglomeration method: Ward’s method. The linkage distance is the measure of similarity between cell subsets, with increasing distance representing decreasing degrees of similarity between samples. (M) GO heatmap and clustering dendrogram with GO terms along the horizontal axis. The heatmap was generated across all cell subsets. P values were calculated by the hypergeometric test and denoted by -log_10_P; overrepresented GO terms were shown in each cell subsets and colored by red and blue in Extend Figure-4M; distance method: correlation; agglomeration method: Ward’s method.

**Table 4.**
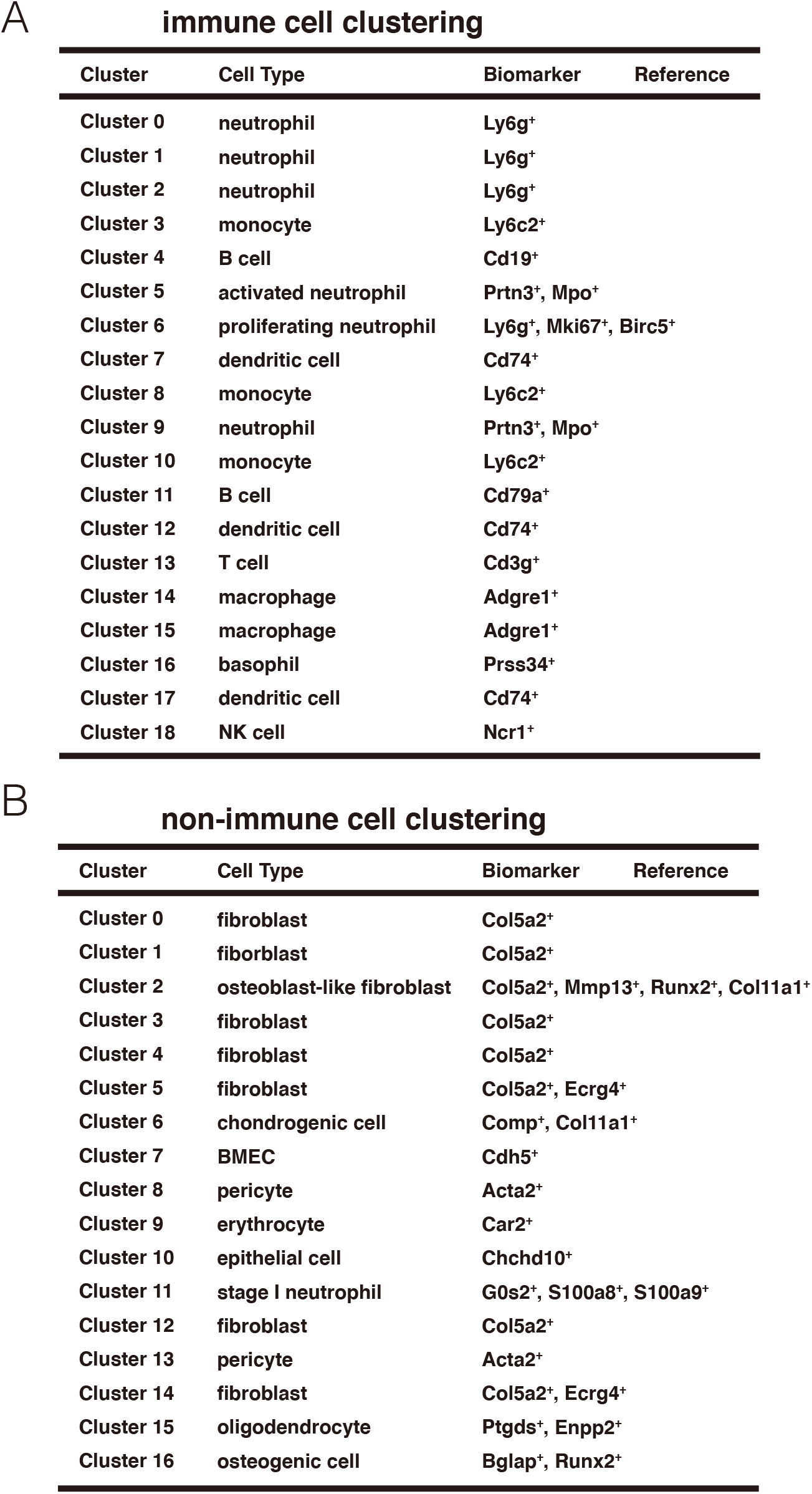
Cell clustering in Fig4/FigS4

**immune cell clustering**
| Cluster | Cell Type | Biomarker | Reference |
| --- | --- | --- | --- |
| Cluster 0 | neutrophil | Ly6g<sup>+</sup> |  |
| Cluster 1 | neutrophil | Ly6g<sup>+</sup> |  |
| Cluster 2 | neutrophil | Ly6g<sup>+</sup> |  |
| Cluster 3 | monocyte | Ly6c2<sup>+</sup> |  |
| Cluster 4 | B cell | Cd19<sup>+</sup> |  |
| Cluster 5 | activated neutrophil | Prtn3<sup>+</sup>, Mpo<sup>+</sup> |  |
| Cluster 6 | proliferating neutrophil | Ly6g<sup>+</sup>, Mki67<sup>+</sup>, Birc5<sup>+</sup> |  |
| Cluster 7 | dendritic cell | Cd74<sup>+</sup> |  |
| Cluster 8 | monocyte | Ly6c2<sup>+</sup> |  |
| Cluster 9 | neutrophil | Prtn3<sup>+</sup>, Mpo<sup>+</sup> |  |
| Cluster 10 | monocyte | Ly6c2<sup>+</sup> |  |
| Cluster 11 | B cell | Cd79a<sup>+</sup> |  |
| Cluster 12 | dendritic cell | Cd74<sup>+</sup> |  |
| Cluster 13 | T cell | Cd3g<sup>+</sup> |  |
| Cluster 14 | macrophage | Adgre1<sup>+</sup> |  |
| Cluster 15 | macrophage | Adgre1<sup>+</sup> |  |
| Cluster 16 | basophil | Prss34<sup>+</sup> |  |
| Cluster 17 | dendritic cell | Cd74<sup>+</sup> |  |
| Cluster 18 | NK cell | Ncr1<sup>+</sup> |  |

**non-immune cell clustering**
| Cluster | Cell Type | Biomarker | Reference |
| --- | --- | --- | --- |
| Cluster 0 | fibroblast | Col5a2<sup>+</sup> |  |
| Cluster 1 | fiborblast | Col5a2<sup>+</sup> |  |
| Cluster 2 | osteoblast-like fibroblast | Col5a2<sup>+</sup>, Mmp13<sup>+</sup>, Runx2<sup>+</sup>, Col11a1<sup>+</sup> |  |
| Cluster 3 | fibroblast | Col5a2<sup>+</sup> |  |
| Cluster 4 | fibroblast | Col5a2<sup>+</sup> |  |
| Cluster 5 | fibroblast | Col5a2<sup>+</sup>, Ecrg4<sup>+</sup> |  |
| Cluster 6 | chondrogenic cell | Comp<sup>+</sup>, Col11a1<sup>+</sup> |  |
| Cluster 7 | BMEC | Cdh5<sup>+</sup> |  |
| Cluster 8 | pericyte | Acta2<sup>+</sup> |  |
| Cluster 9 | erythrocyte | Car2<sup>+</sup> |  |
| Cluster 10 | epithelial cell | Chchd10<sup>+</sup> |  |
| Cluster 11 | stage I neutrophil | G0s2<sup>+</sup>, S100a8<sup>+</sup>, S100a9<sup>+</sup> |  |
| Cluster 12 | fibroblast | Col5a2<sup>+</sup> |  |
| Cluster 13 | pericyte | Acta2<sup>+</sup> |  |
| Cluster 14 | fibroblast | Col5a2<sup>+</sup>, Ecrg4<sup>+</sup> |  |
| Cluster 15 | oligodendrocyte | Ptgds<sup>+</sup>, Enpp2<sup>+</sup> |  |
| Cluster 16 | osteogenic cell | Bglap<sup>+</sup>, Runx2<sup>+</sup> |  |

Next, WGCNA identified seven modules associated with phenotype and clustering respectively (MEgreen, MEred and MEturquoise were negatively associated with phenotype and clustering, MEblack is positively correlated with phenotype, MEpink is positively correlated with clustering, MEmagenta and MEgrey are positively correlated with phenotype and clustering, Figure S4B-D). Genes in each module were distributed in cluster 1, 6, 9, 10 (MEgreen); cluster 1, 2, 3, 10 (MEred); cluster 4, 11 (MEblack); cluster 13, 18 (MEmagenta); cluster 14, 15 (MEpink); all cluster (MEturquoise and MEgrey) (Figure S3E), indicating that neutrophils worked in the whole repair process, and B cells and Basophils play important roles in bone remodeling.

In the non-immune cell group, we identified 17 clusters (Figure 4H) and the inconsistency between n21 and sc21 via PSA and DMA, respectively (Figures 4I and S4F). Cluster similarities were performed by relatedness of CGAA and functional enrichment analysis (Figure 4J-L), showing that cluster 10-11 (cluster 10 mainly in n21, cluster 11 in both group), cluster 8-13 (cluster 8 mainly in sc21, cluster 13 mainly in n21), cluster 2-6 (both mainly in sc21), cluster 5-14 (both mainly in n21) have similarity in function. The detailed cell identities and corresponding function could be found in Table 4B, Figure 4G and Extend file for Fig 4G. Then, we used WGCNA to look for the difference of non-immune cells between sc21 and n21 group (Figures S4G and S4H), and found that four modules were associated with phenotype and clustering, respectively (MEgreen was positively correlated with phenotype and negatively correlated with clustering, MEyellow was positively correlated with clustering, MEpink and MEblack are negatively correlated with phenotype and positively correlated with clustering, Figure S4I). By extracting genes in each module, we found that they were mainly in, cluster 0, 1, 11, 12 (MEgreen); cluster 5, 7, 10, 13, 14, 15, 16 (MEpink); cluster 8, 13 (MEyellow); cluster 10, 11 (MEblack) (Figure S4J), illustrating Ecrg4^+^fibroblast, Acta2^+^ pericyte, Chchd10^+^ epithelial cells and Cdh5^+^ endothelial cells contributing to the whole repair process.

### Roles of innate and adaptive immune cells during the progression of bone repair

In order to test whether the process of bone repair would be affected by immune reversion after the occurrence of immune deficiency, we introduced the hole drilling model into NPSG mice humanized with CD34^+^ cells, which can distinguish the bone-healing activity between myeloid and lymphoid cells, since bone defects in mice could be regenerated by crosstalk of both human and mouse cells (*20*). We obtained tissues in bone repair site at day 7, 14, and 21, and then, tissues were subjected to single-cell sequencing (Figure 5A).

**Figure 5.**
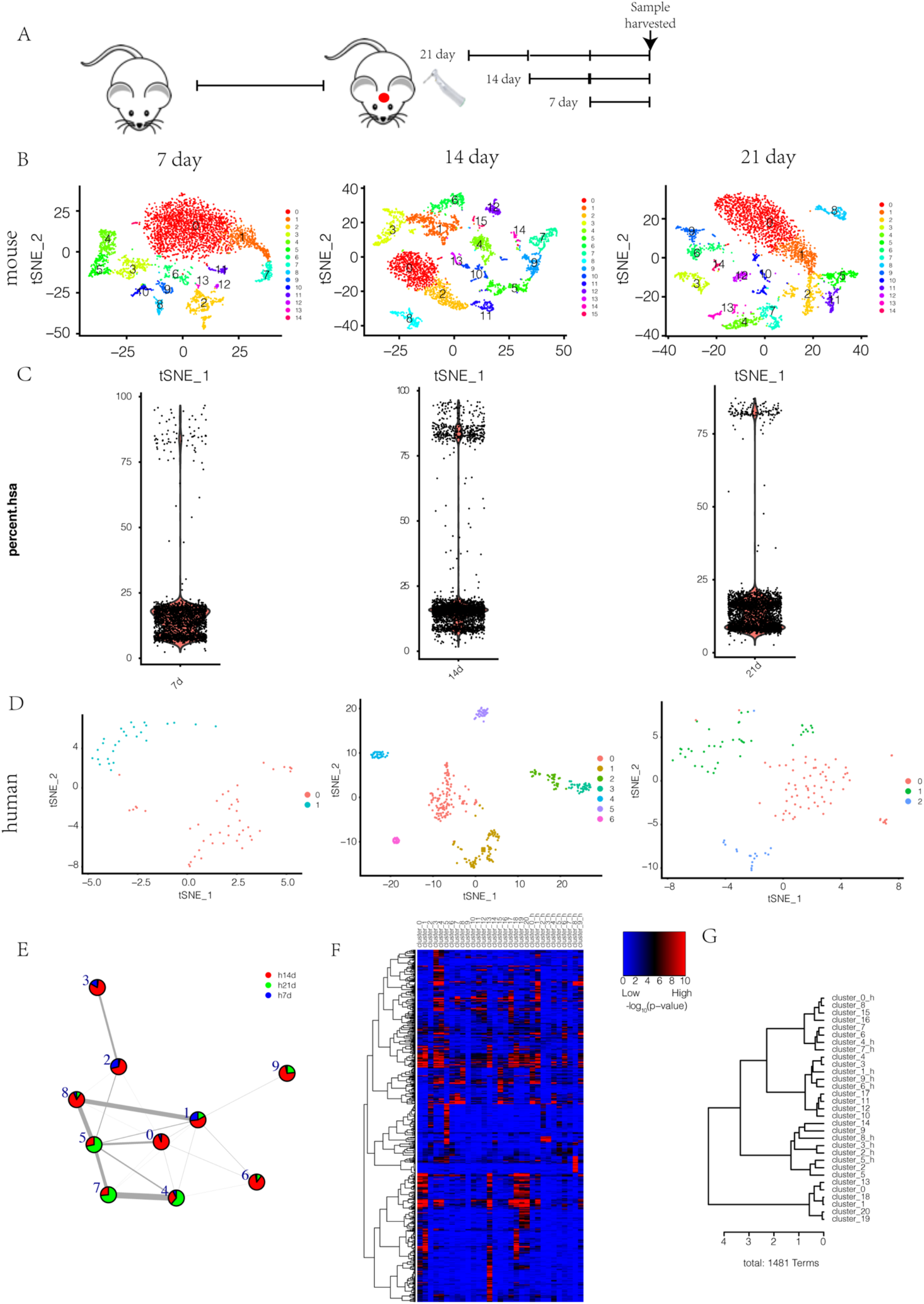
Innate immune cells play a compensatory role during the progression of bone repair in absence of adaptive immune cells. (A) Study design of bone repair in humanized mouse. (B) Cell census at bone reformation site colored by cell subsets from t-SNE based on mouse genome. (C) Distribution of mouse cells and human cells at post-surgery time points. (D) Cell census at bone reformation site colored by cell subsets from t-SNE based on human genome. (E) Relatedness (edges, width indicates strength) between clusters (nodes, colored as post-surgery time points) based on cluster graph abstraction. (F) GO heatmap and clustering dendrogram of human cells and mouse cells with GO terms along the horizontal axis. The heatmap was generated across all cell subsets. P values were calculated by the hypergeometric test and denoted by -log_10_P; overrepresented GO terms were shown in each cell subsets and colored by red and blue in Extend Figure-5F; distance method: correlation; agglomeration method: Ward’s method. (G) Cluster dendrogram of cell subsets in F based on the patterns of GO enrichment; distance method: correlation; agglomeration method: Ward’s method. The linkage distance is the measure of similarity between cell subsets, with increasing distance representing decreasing degrees of similarity between samples.

The corresponding cell populations in the bone injury site of humanized mice were clustered by t-SNE method (4641, 3872, 3620 cells, respectively, Figure 5B). At the same time, since we traced back human CD34^+^ cells into mice, we also used the human genome as a reference to map the data, successfully separated a number of human cells, and performed the corresponding clustering (Figure 5C-D).

After merging and analyzing the human basal cell populations at the three time points, we got 16 clusters (Figure S5A). Using both PSA and DMA, we found the appearance of cell state progression over time (Figures S5B-C). After obtaining the marker gene expression map (Figure S5D), we performed relatedness analysis based on cluster graph abstraction, and found that the clusters containing cells in day 7 and day 14 were closer in function, and those containing cells in day 14 and 21 were functionally close. Annotation of those human-based cell populations at different time points revealed that at day 7 and 14, T cells, dendritic cells, and stem cells were mainly distributed, and at day 21, B cells were the main types at that time (Figure 5E and Table 5A).

**Table 5.**
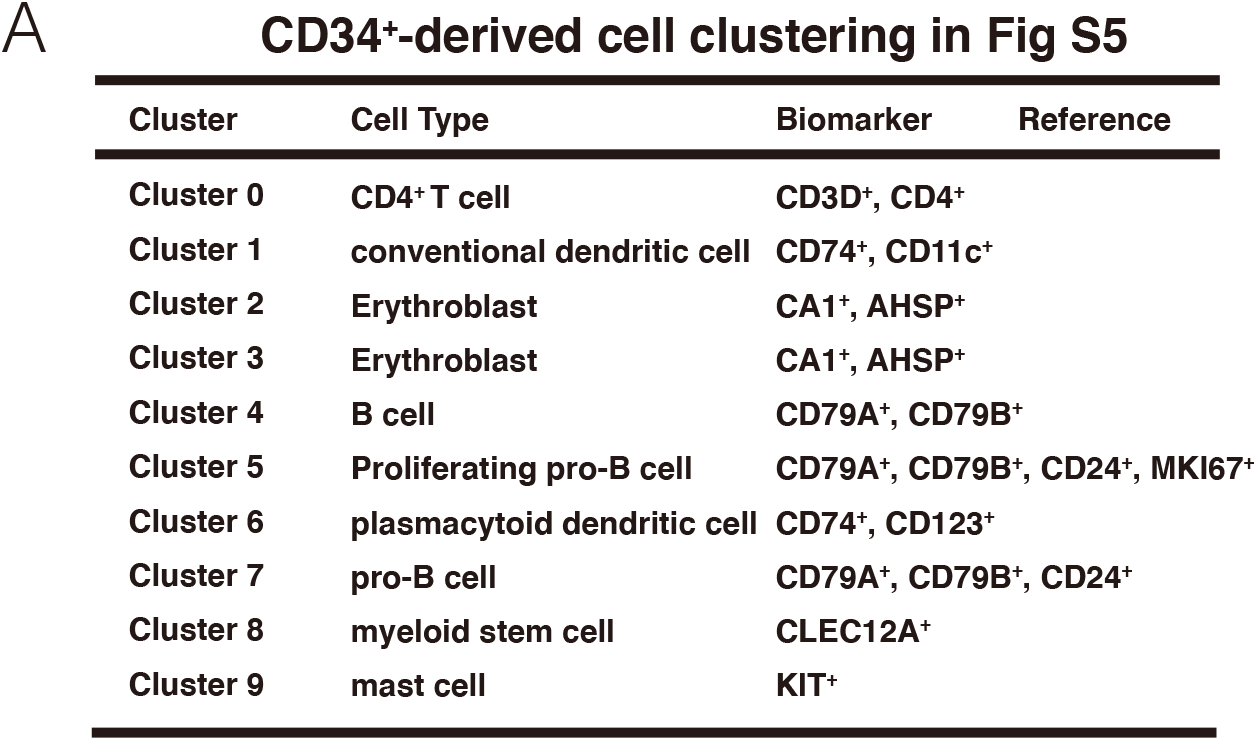
CD34^+^-derived cell clustering in Fig S5

| CD34<sup>+</sup>-derived cell clustering in Fig S5 |  |  |  |
| --- | --- | --- | --- |
| Cluster | Cell Type | Biomarker | Reference |
| Cluster 0 | CD4<sup>+</sup>T cell | CD3D<sup>+</sup>, CD4<sup>+</sup> |  |
| Cluster 1 | conventional dendritic cell | CD74<sup>+</sup>, CD11c<sup>+</sup> |  |
| Cluster 2 | Erythroblast | CA1<sup>+</sup>, AHSP<sup>+</sup> |  |
| Cluster 3 | Erythroblast | CA1<sup>+</sup>, AHSP<sup>+</sup> |  |
| Cluster 4 | B cell | CD79A<sup>+</sup>, CD79B<sup>+</sup> |  |
| Cluster 5 | Proliferating pro-B cell | CD79A<sup>+</sup>, CD79B<sup>+</sup>, CD24<sup>+</sup>, MKI67<sup>+</sup> |  |
| Cluster 6 | plasmacytoid dendritic cell | CD74<sup>+</sup>, CD123<sup>+</sup> |  |
| Cluster 7 | pro-B cell | CD79A<sup>+</sup>, CD79B<sup>+</sup>, CD24<sup>+</sup> |  |
| Cluster 8 | myeloid stem cell | CLEC12A<sup>+</sup> |  |
| Cluster 9 | mast cell | KIT<sup>+</sup> |  |

We then compared the immune cells of normal ICR mice at day 7, 14, and 21 with these human basal cells by GO-based gene enrichment analysis, and illustrated that human cluster 0-mouse cluster 8; human cluster 4, 7-mouse cluster 6,7; human cluster 1,6,9-mouse cluster 3,4; human cluster 2, 3, 5, 8-mouse cluster 2, 5, 9, 14 were functionally close (Figure 5F and G).

### Integrated cellular mapping and hub genes in the different stage of bone repair

In order to clarify whether there were differences in given types of cells at different times at the bone repair site after immune deficiency and immune recovery, we compared the cells of ICR mice (sc), NPSG mice (n) and CD34 positive mice (cd) at the same time, horizontally.

At day 7, the immune cells of bone repair tissues from the three kinds of mice were divided into 18 clusters (Figure 6A). PSA showed that the types of immune cells in the sc7 group were significantly different from the n7 and cd7 groups (Figure 6B). By CGAA (Figures 6C) and pairwise similarity analysis based on functional enrichment analysis results (Figure 6D), it was found that cluster 11-14 (cluster 11 mainly in sc7, cluster 14 mainly in sc7 and n7) and cluster 4-12 (cluster 4 mainly in n7 and cd7; cluster 12 mainly in cd7) were more similar in gene expression and function. Other clusters were identified in Table 6A, and their functions are reflected in Figure 6E and Extend file for Fig 6E). By listing genes involved in the main metabolic pathways, we found that at day 7, neutrophils (cluster 0,2,3,7) maintained a low level of metabolism, and M2 macrophage (cluster 5, 9) prefer fatty acid oxidation (FAO), but not glycolysis, while osteoclasts (cluster 10) have a high level of all the metabolism pathways (Figure 6F).

**Figure 6.**
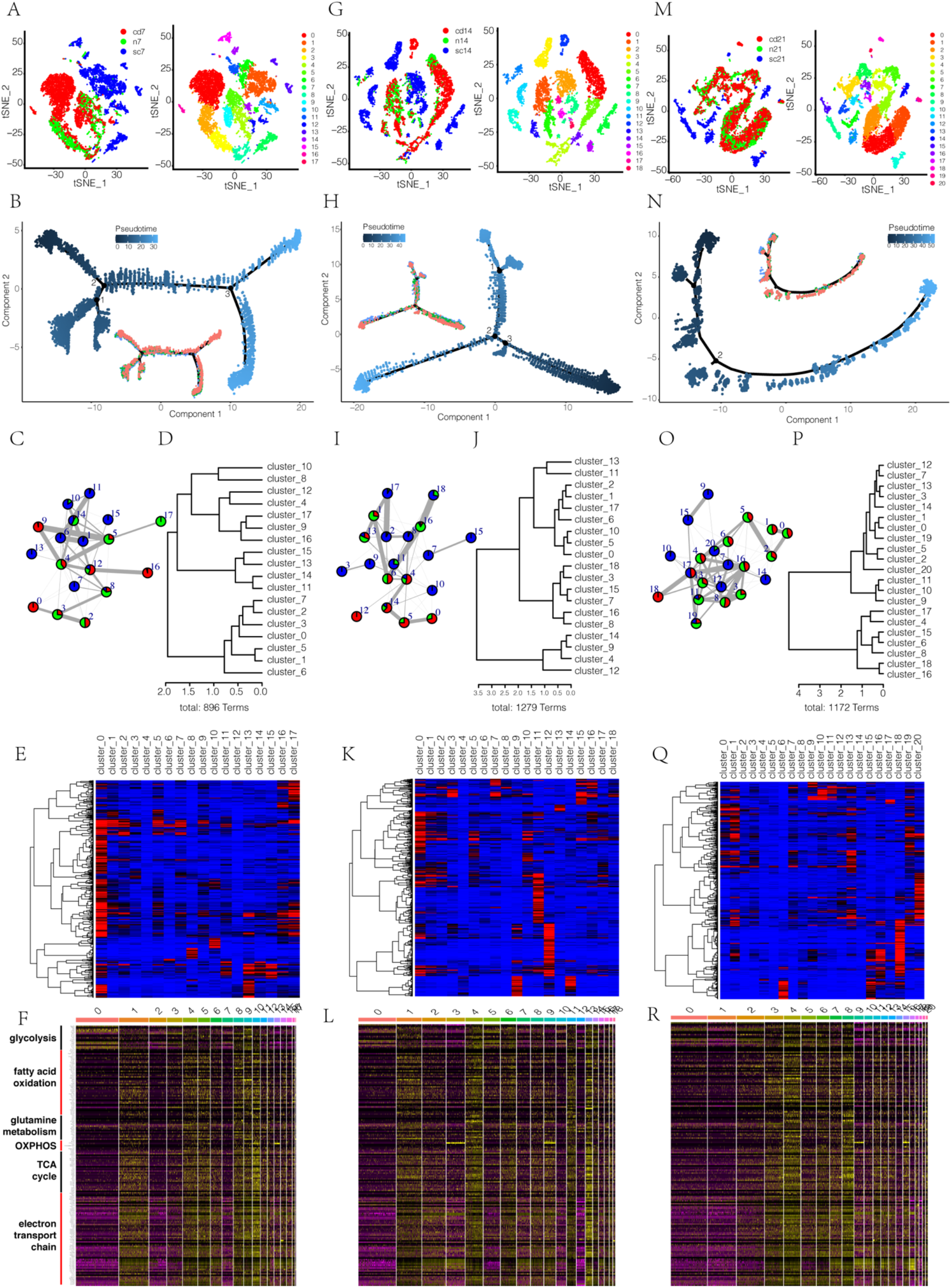
Hub genes and cell types in different stage of bone repair. (A) Cell census at bone reformation site colored by cell subsets and different mouse models at post-surgery 7 day from t-SNE and diffusion map analysis. (A) (B) Pseudotime analysis of cells in A colored by pseudotime and mouse models. (C) Relatedness (edges, width indicates strength) between clusters (nodes, colored as mouse models) based on cluster graph abstraction. (D) Cluster dendrogram of cell subsets in C based on the patterns of GO enrichment; distance method: correlation; agglomeration method: Ward’s method. The linkage distance is the measure of similarity between cell subsets, with increasing distance representing decreasing degrees of similarity between samples. (E) GO heatmap and clustering dendrogram of cell subsets in A with GO terms along the horizontal axis. The heatmap was generated across all cell subsets. P values were calculated by the hypergeometric test and denoted by -log_10_P; overrepresented GO terms were shown in each cell subsets and colored by red and blue in Extend Figure-6E; distance method: correlation; agglomeration method: Ward’s method. (F) Marker genes related to metabolisms in different cell subsets in A. (G) Cell census at bone reformation site colored by cell subsets and different mouse models at post-surgery 14 day from t-SNE and diffusion map analysis. (H) Pseudotime analysis of cells in G colored by pseudotime and mouse models. (I) Relatedness (edges, width indicates strength) between clusters (nodes, colored as mouse models) based on cluster graph abstraction. (J) Cluster dendrogram of cell subsets in I based on the patterns of GO enrichment; distance method: correlation; agglomeration method: Ward’s method. The linkage distance is the measure of similarity between cell subsets, with increasing distance representing decreasing degrees of similarity between samples. (K) GO heatmap and clustering dendrogram of cell subsets in G with GO terms along the horizontal axis. The heatmap was generated across all cell subsets. P values were calculated by the hypergeometric test and denoted by -log_10_P; overrepresented GO terms were shown in each cell subsets and colored by red and blue in Extend Figure-6K; distance method: correlation; agglomeration method: Ward’s method. (L) Marker genes related to metabolisms in different cell subsets in G. (M) Cell census at bone reformation site colored by cell subsets and different mouse models at post-surgery 21 day from t-SNE and diffusion map analysis. (N) Pseudotime analysis of cells in M colored by pseudotime and mouse models. (O) Relatedness (edges, width indicates strength) between clusters (nodes, colored as mouse models) based on cluster graph abstraction. (P) Cluster dendrogram of cell subsets in M based on the patterns of GO enrichment; distance method: correlation; agglomeration method: Ward’s method. The linkage distance is the measure of similarity between cell subsets, with increasing distance representing decreasing degrees of similarity between samples. (Q) GO heatmap and clustering dendrogram of cell subsets in M with GO terms along the horizontal axis. The heatmap was generated across all cell subsets. P values were calculated by the hypergeometric test and denoted by -log_10_P; overrepresented GO terms were shown in each cell subsets and colored by red and blue in Extend Figure-6Q; distance method: correlation; agglomeration method: Ward’s method. (R) Marker genes related to metabolisms in different cell subsets in M.

**Table 6.** Cell clustering in Fig 6A/S6A

| A<br>immune cell clustering in Fig 6A |  |  |  |
| --- | --- | --- | --- |
| Cluster | Cell Type | Biomarker | Reference |
| Cluster 0 | neutrophil | Csf3r<sup>+</sup> |  |
| Cluster 1 | Microglial cell | Cx3cr1<sup>+</sup>, Aif1<sup>+</sup> |  |
| Cluster 2 | neutrophil | Csf3r<sup>+</sup>, Ly6g<sup>+</sup> |  |
| Cluster 3 | neutrophil | Csf3r<sup>+</sup>, Ly6g<sup>+</sup> |  |
| Cluster 4 | monocyte | Ly6c2<sup>+</sup>, Lyz2<sup>+</sup>, Ccr2<sup>+</sup> |  |
| Cluster 5 | M2, macrophage | F4/80<sup>+</sup>, Mrc1<sup>+</sup> |  |
| Cluster 6 | monocyte | Ly6c2<sup>+</sup>, Lyz2<sup>+</sup>, Ccr2<sup>+</sup> |  |
| Cluster 7 | neutrophil | Csf3r<sup>+</sup>, Ly6g<sup>+</sup> |  |
| Cluster 8 | myeloid stem cell | Clec12a<sup>+</sup> |  |
| Cluster 9 | M2, macrophage | F4/80<sup>+</sup>, Arg1<sup>+</sup> |  |
| Cluster 10 | osteoclast | Ctsk<sup>+</sup> |  |
| Cluster 11 | dendritic cell | Cd74<sup>+</sup> |  |
| Cluster 12 | fibroblast | Col5a2<sup>+</sup> |  |
| Cluster 13 | B cell | Cd79a<sup>+</sup>, Cd79b<sup>+</sup> |  |
| Cluster 14 | plasmacytoid dendritic cell | Siglech<sup>+</sup>, Bst2<sup>+</sup> |  |
| Cluster 15 | T cell | Cd3g<sup>+</sup> |  |
| Cluster 16 | stem cell | Ctla2a<sup>+</sup> |  |
| Cluster 17 | Microglial cell | Cx3cr1<sup>+</sup>, Aif1<sup>+</sup> |  |

| B<br>non-immune cell clustering in Fig S6A |  |  |  |
| --- | --- | --- | --- |
| Cluster | Cell Type | Biomarker | Reference |
| Cluster 0 | pericyte | Acta2<sup>+</sup>, Ptprd<sup>+</sup> |  |
| Cluster 1 | fibroblast | Col5a2<sup>+</sup>, Ecrg4<sup>+</sup> |  |
| Cluster 2 | endothelial cell | Ly6c1<sup>+</sup>, Pi16<sup>+</sup> |  |
| Cluster 3 | fibroblast | Col5a2<sup>+</sup> |  |
| Cluster 4 | fibroblast | Col5a2<sup>+</sup> |  |
| Cluster 5 | pericyte | Acta2<sup>+</sup> |  |
| Cluster 6 | endothelial cell | Pecam1<sup>+</sup>, Cdh5<sup>+</sup> |  |
| Cluster 7 | smooth muscle cell | Rgs5<sup>+</sup> |  |
| Cluster 8 | pericyte cell, proliferating | Acta2<sup>+</sup>, Mki67<sup>+</sup> |  |
| Cluster 9 | fibroblast | Col5a2<sup>+</sup>, Ecrg4<sup>+</sup> |  |
| Cluster 10 | erythrocytes | Car2<sup>+</sup> |  |
| Cluster 11 | chondrogenic cell | Comp<sup>+</sup>, Acan<sup>+</sup> |  |
| Cluster 12 | endothelial cell | Pecam1<sup>+</sup>, Cdh5<sup>+</sup> |  |
| Cluster 13 | microglial cell | Cx3cr1<sup>+</sup>, Ctss<sup>+</sup> |  |
| Cluster 14 | monocyte | Cd14<sup>+</sup>, Il1b<sup>+</sup> |  |
| Cluster 15 | oligodendrocyte | Mbp<sup>+</sup>, Ptgsd<sup>+</sup> |  |
| Cluster 16 | smooth muscle cell | Rgs5<sup>+</sup> |  |
| Cluster 17 | fibroblast | Col5a2<sup>+</sup> |  |
| Cluster 18 | osteogenic cell | Bglap<sup>+</sup>, Sp7<sup>+</sup> |  |
| Cluster 19 | oligodendrocyte | Mbp<sup>+</sup>, Ptgsd<sup>+</sup> |  |
| Cluster 20 | oligodendrocyte | Mbp<sup>+</sup>, Mal<sup>+</sup> |  |

On the non-immune cell side, we identified 21 clusters (Figure S6A). PSA showed that the sc group was significantly different from the n and cd groups in non-immune cell types (Figure S6B). CGAA and functionally interblock similarity analysis identified that clusters 11-18, 7-16, 6-12, 15-19-20 and 13-14 have closer relations in the gene expression profile and GO-based function (Figure S6C-D). The detailed cell identities and functions are reflected in Table 6B, Figure S6E and Extend file for Fig S6E). According to genes related to the main metabolic pathways, angiogenesis and ossification, we found that at day 7, pericyte (cluster 5) owns a low metabolic level, while osteogenic cells (cluster 18) prefer fatty acid oxidation, but not glycolysis (Figure S6F). Cdh5^+^ endothelial cells (cluster 12) contribute to angiogenesis, while osteogenic gene were distributed in pericyte (cluster 0), Ecrg4^+^fibroblast (cluster 1, 9), Pi16^+^ endothelial cells (cluster 2), chondrogenic cells (cluster 11) and osteogenic cells (cluster 18) (Figure S6F). Likewise, we analyzed 14-day and 21-day data in the same manner. According to the 14-day immune cells, we identified 19 clusters (Figure 6G), similarly, sc group was significantly different from n and cd groups (Figure 6H). Cluster 1-2-17, 7-15, 8-16 are relatively close in gene expression and function (Figures 6I and 6J). The detailed information is reflected in Table 7A, Figure 6K and Extend file for Fig 6K. From Figure 6L, M2, macrophage and microglial cells (cluster 2,3) prefers FAO, while the metabolic levels in myeloid cells (eosinophil in cluster 4, neutrophils in cluster 5,10,14, myeloid-derived suppressor cells [MDSCs] in cluster 6) and osteoclast (cluster 13) were high (Figure 6L). The 14-day non-immune cells were divided into 21 clusters (Figure S6G) and different between sc group and the other two groups (Figure S6H). Clusters 17-19, 11-16, 10-12 and 2-8 are functionally close (Figures S6I and S6J). The detailed information is shown in Table 7B, Figure S6K and Extend file for Fig S6K. The metabolic gene profile showed that epithelial cell in cluster 14 preferred glucose to be energy resource, while osteogenic cell in cluster 15 used fatty acid (Figure S6L). Endothelial cells (cluster 4, 7) and fibroblasts (cluster 2, 8), smooth muscle cells (cluster 10) contribute to angiogenesis, while fibroblasts (cluster 3,4,), chondrogenic cell (cluster 6), osteogenic cell (cluster 15) are related to ossification (Figure S6L).

**Table 7.**
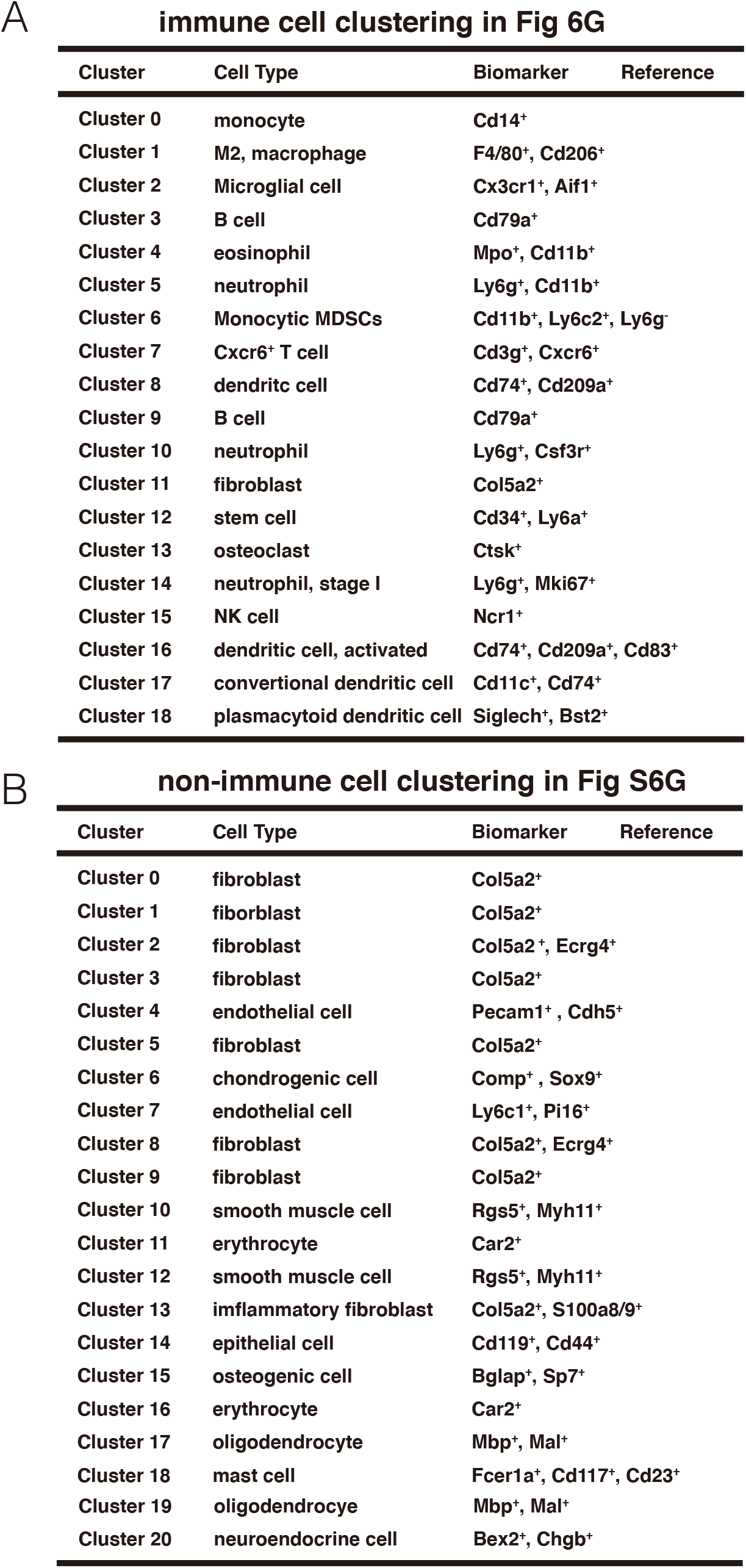
Cell clustering in Fig6G/S6G

**immune cell clustering in Fig 6G**
| Cluster | Cell Type | Biomarker | Reference |
| --- | --- | --- | --- |
| Cluster 0 | monocyte | Cd14<sup>+</sup> |  |
| Cluster 1 | M2, macrophage | F4/80<sup>+</sup>, Cd206<sup>+</sup> |  |
| Cluster 2 | Microglial cell | Cx3cr1<sup>+</sup>, Aif1<sup>+</sup> |  |
| Cluster 3 | B cell | Cd79a<sup>+</sup> |  |
| Cluster 4 | eosinophil | Mpo<sup>+</sup>, Cd11b<sup>+</sup> |  |
| Cluster 5 | neutrophil | Ly6g<sup>+</sup>, Cd11b<sup>+</sup> |  |
| Cluster 6 | Monocytic MDSCs | Cd11b<sup>+</sup>, Ly6c2<sup>+</sup>, Ly6g<sup>-</sup> |  |
| Cluster 7 | Cxcr6<sup>+</sup> T cell | Cd3g<sup>+</sup>, Cxcr6<sup>+</sup> |  |
| Cluster 8 | dendritic cell | Cd74<sup>+</sup>, Cd209a<sup>+</sup> |  |
| Cluster 9 | B cell | Cd79a<sup>+</sup> |  |
| Cluster 10 | neutrophil | Ly6g<sup>+</sup>, Csf3r<sup>+</sup> |  |
| Cluster 11 | fibroblast | Col5a2<sup>+</sup> |  |
| Cluster 12 | stem cell | Cd34<sup>+</sup>, Ly6a<sup>+</sup> |  |
| Cluster 13 | osteoclast | Ctsk<sup>+</sup> |  |
| Cluster 14 | neutrophil, stage I | Ly6g<sup>+</sup>, Mki67<sup>+</sup> |  |
| Cluster 15 | NK cell | Ncr1<sup>+</sup> |  |
| Cluster 16 | dendritic cell, activated | Cd74<sup>+</sup>, Cd209a<sup>+</sup>, Cd83<sup>+</sup> |  |
| Cluster 17 | conversional dendritic cell | Cd11c<sup>+</sup>, Cd74<sup>+</sup> |  |
| Cluster 18 | plasmacytoid dendritic cell | Siglech<sup>+</sup>, Bst2<sup>+</sup> |  |

**non-immune cell clustering in Fig S6G**
| Cluster | Cell Type | Biomarker | Reference |
| --- | --- | --- | --- |
| Cluster 0 | fibroblast | Col5a2<sup>+</sup> |  |
| Cluster 1 | fibroblast | Col5a2<sup>+</sup> |  |
| Cluster 2 | fibroblast | Col5a2<sup>+</sup>, Ecrg4<sup>+</sup> |  |
| Cluster 3 | fibroblast | Col5a2<sup>+</sup> |  |
| Cluster 4 | endothelial cell | Pecam1<sup>+</sup>, Cdh5<sup>+</sup> |  |
| Cluster 5 | fibroblast | Col5a2<sup>+</sup> |  |
| Cluster 6 | chondrogenic cell | Comp<sup>+</sup>, Sox9<sup>+</sup> |  |
| Cluster 7 | endothelial cell | Ly6c1<sup>+</sup>, Pi16<sup>+</sup> |  |
| Cluster 8 | fibroblast | Col5a2<sup>+</sup>, Ecrg4<sup>+</sup> |  |
| Cluster 9 | fibroblast | Col5a2<sup>+</sup> |  |
| Cluster 10 | smooth muscle cell | Rgs5<sup>+</sup>, Myh11<sup>+</sup> |  |
| Cluster 11 | erythrocyte | Car2<sup>+</sup> |  |
| Cluster 12 | smooth muscle cell | Rgs5<sup>+</sup>, Myh11<sup>+</sup> |  |
| Cluster 13 | inflammatory fibroblast | Col5a2<sup>+</sup>, S100a8/9<sup>+</sup> |  |
| Cluster 14 | epithelial cell | Cd119<sup>+</sup>, Cd44<sup>+</sup> |  |
| Cluster 15 | osteogenic cell | Bglap<sup>+</sup>, Sp7<sup>+</sup> |  |
| Cluster 16 | erythrocyte | Car2<sup>+</sup> |  |
| Cluster 17 | oligodendrocyte | Mbp<sup>+</sup>, Mal<sup>+</sup> |  |
| Cluster 18 | mast cell | Fcer1a<sup>+</sup>, Cd117<sup>+</sup>, Cd23<sup>+</sup> |  |
| Cluster 19 | oligodendrocyte | Mbp<sup>+</sup>, Mal<sup>+</sup> |  |
| Cluster 20 | neuroendocrine cell | Bex2<sup>+</sup>, Chgb<sup>+</sup> |  |

In the 21-day immune cells, we confirmed 21 clusters (Figure 6M) and the difference among the three groups (Figure 6N). Cluster 7-12, 0-1, 2-5 and 4-17 have close relatedness in terms of gene expression and function (Figures 6O and 6P). The detailed contents are showed in Table 8A, Figure 6Q and Extend file for Fig 6Q. Neutrophils in cluster 0,1,2, and monocyte in cluster 14 are in a low energy-producing state, while monocyte (cluster 3, 7, 8) and neutrophils (cluster 5,6) have a high expression of energy-producing genes (Figure 6R). The 21-day non-immune cells were divided into 20 clusters (Figure S6M-N). Clusters 7-11, 1-4, 6-18, 14 -15, are relatively close in terms of gene expression and function (Figures S6O and S6P). The detailed contents are showed in Table 8B, Figure S6Q and Extend file for Fig S6Q. Endothelial cells in cluster 0 and stem cell in cluster 14 prefer to use glutamine metabolism. Endothelial cells in cluster 0, 1, 7, 11 contributed to angiogenesis, and fibroblasts (cluster 2, 3, 4, 9, 15), chondrogenic cells (cluster 5) and osteogenic cells (cluster 19) have a higher expression of ossification-related genes (Figure S6R).

**Table 8.** Cell clustering in Fig 6M/S6M

**immune cell clustering in Fig 6M**
| Cluster | Cell Type | Biomarker | Reference |
| --- | --- | --- | --- |
| Cluster 0 | neutrophil | Ly6g<sup>+</sup>, Csf3r<sup>+</sup> |  |
| Cluster 1 | neutrophil | Ly6g<sup>+</sup>, Csf3r<sup>+</sup> |  |
| Cluster 2 | neutrophil, stage I | Ly6g<sup>+</sup>, Camp<sup>+</sup> |  |
| Cluster 3 | monocyte | Cd11b<sup>+</sup>, Ly6c2<sup>+</sup> |  |
| Cluster 4 | stem cell | Cd44<sup>+</sup>, Ctla2a<sup>+</sup> |  |
| Cluster 5 | neutrophil, stage I | Ly6g<sup>+</sup>, Camp<sup>+</sup> |  |
| Cluster 6 | neutrophil, proliferating | Ly6g<sup>+</sup>, Mki67<sup>+</sup> |  |
| Cluster 7 | monocyte | Cd11b<sup>+</sup>, Csf1r<sup>+</sup>, S100a4<sup>+</sup> |  |
| Cluster 8 | monocyte | Cd11b<sup>+</sup>, Ly6c2<sup>+</sup> |  |
| Cluster 9 | B cell | Cd79a<sup>+</sup>, Cd79b<sup>+</sup> |  |
| Cluster 10 | Cxcr6<sup>+</sup> T cell | Cd3g<sup>+</sup>, Cxcr6<sup>+</sup> |  |
| Cluster 11 | plasmacytoid dendritic cell | Siglech<sup>+</sup>, Bst2<sup>+</sup> |  |
| Cluster 12 | dendritic cell | Cd74<sup>+</sup>, Cd209a<sup>+</sup> |  |
| Cluster 13 | microglial cell | Cx3cr1<sup>+</sup> |  |
| Cluster 14 | monocyte | Cd11b<sup>+</sup>, Ly6c2<sup>+</sup> |  |
| Cluster 15 | B cell | Cd79a<sup>+</sup>, Cd79b<sup>+</sup> |  |
| Cluster 16 | stem cell | Cd44<sup>+</sup> |  |
| Cluster 17 | erythrocyte | Car2<sup>+</sup> |  |
| Cluster 18 | B cell progenitor | Sox4<sup>+</sup> |  |
| Cluster 19 | basophil | Prss34<sup>+</sup> |  |
| Cluster 20 | osteoclast | Ctsk<sup>+</sup> |  |

**non-immune cell clustering in Fig S6M**
| Cluster | Cell Type | Biomarker | Reference |
| --- | --- | --- | --- |
| Cluster 0 | endothelial cell | Ly6c1<sup>+</sup>, Pi16<sup>+</sup> |  |
| Cluster 1 | endothelial cell | Ly6c1<sup>+</sup>, Pi16<sup>+</sup> |  |
| Cluster 2 | fibroblast | Col5a2<sup>+</sup> |  |
| Cluster 3 | fibroblast | Col5a2<sup>+</sup> |  |
| Cluster 4 | fibroblast | Col5a2<sup>+</sup> |  |
| Cluster 5 | chondrogenic cell | Comp<sup>+</sup>, Runx2<sup>+</sup> |  |
| Cluster 6 | fibroblast | Col5a2<sup>+</sup>, Ecrg4<sup>+</sup> |  |
| Cluster 7 | endothelial cell | Pecam1<sup>+</sup>, Cdh5<sup>+</sup> |  |
| Cluster 8 | pericyte | Acta2<sup>+</sup> |  |
| Cluster 9 | fibroblast | Col5a2<sup>+</sup>, Ecrg4<sup>+</sup> |  |
| Cluster 10 | erythrocyte | Car2<sup>+</sup> |  |
| Cluster 11 | endothelial cell | Pecam1<sup>+</sup>, Cdh5<sup>+</sup> |  |
| Cluster 12 | erythrocyte | Car2<sup>+</sup> |  |
| Cluster 13 | oligodendrocyte | Kcna1<sup>+</sup>, Mbp<sup>+</sup> |  |
| Cluster 14 | hematopoietic stem cell | Ctla2a<sup>+</sup>, c-Kit<sup>+</sup> |  |
| Cluster 15 | inflammatory fibroblast | Col5a2<sup>+</sup>, S100a8/9<sup>+</sup> |  |
| Cluster 16 | pericyte | Acta2<sup>+</sup> |  |
| Cluster 17 | granulocyte | Csf3r<sup>+</sup>, Cd11b<sup>+</sup> |  |
| Cluster 18 | fibroblast | Col5a2<sup>+</sup>, Ecrg4<sup>+</sup> |  |
| Cluster 19 | osteogenic cell | Bglap<sup>+</sup>, Runx2<sup>+</sup> |  |

### Human adaptive immune cells regulate bone repair process in a similar manner with mouse

In order to investigate the hub genes contributing to the bone repair process, we used WGCNA to conduct in-depth analysis of data from bone repair tissue with different phenotypes at different times, looking for the main cell types for phenotype and clustering. At 7 days of immune cell populations, we identified MEturquoise and MRgrey modules, which were strongly correlated with phenotype and clustering, respectively (Figures 7A-C). Part of MEturquoise is mainly in cluster 0, 2, 3, and the other part is mainly in cluster 1, 4, 5, 7, 8, 9. MEgrey is distributed in all clusters (Figure 7D). The corresponding functions of these two MEs are mainly: angiogenesis, T cell activation, and cell death, neutrophil migration, etc. (Figure 7E), while the corresponding human-based cell population is monocyte (h7cluster 0) (Figure 7F). In non-immune cells, we confirmed that MEpurple, MEyellow, and MEturquoise have strong correlations with phenotype and clustering respectively (Figures S7A-C). MEpurple is mainly in cluster 13, 14, MEyellow is cluster 1, 9, 16∼18, and MEturquoise is mainly in cluster 15 (Figures S7D), and its functions are as followed: MEpurple, neutrophil chemical migration, MEturquoise, neurogenesis and adipogenesis, and MEyellow, programmed cell death (Figures S7E).

**Figure 7.**
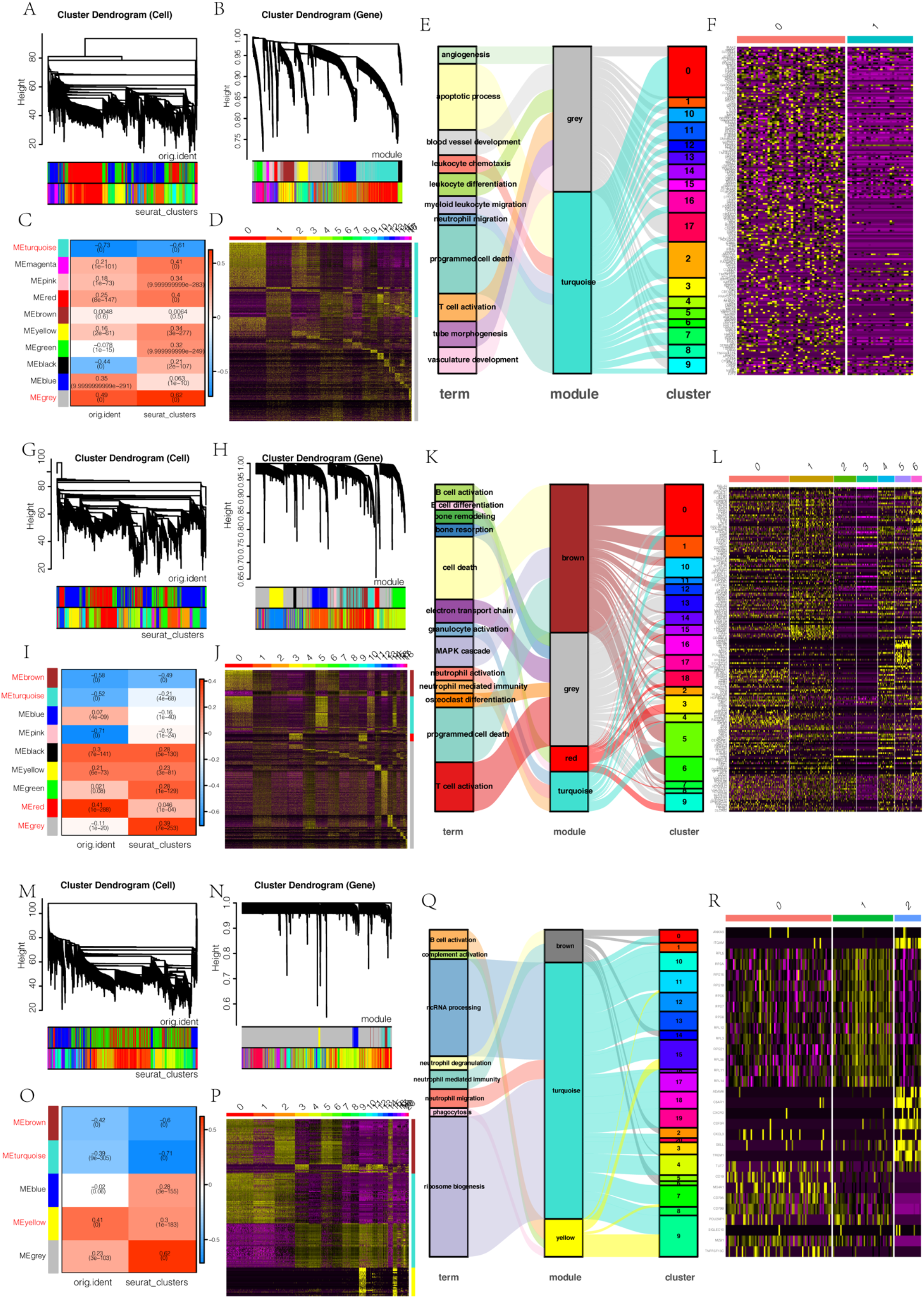
Human adaptive immune cells act similar to mouse in regulating bone repair process. (A) Cell and (B) gene trees based on WCGNA in post-surgery 7 day. (C) Correlations of significant modules to cell subsets in Figure 6A and mouse models. (D) Expressions of genes involved in module turquoise and grey in different cell subsets in Figure 6A. (E) Sankey maps of immune-related GO terms towards modules and cell subsets with corresponding homologous genes to human cells in post-surgery 7 day (F). (G) Cell and (H) gene trees based on WCGNA in post-surgery 14 day. (I) Correlations of significant modules to cell subsets in Figure 6G and mouse models. (J) Expressions of genes involved in module turquoise and grey in different cell subsets in Figure 6G. (K) Sankey maps of immune-related GO terms towards modules and cell subsets with corresponding homologous genes to human cells in post-surgery 14 day (L). (M) Cell and (N) gene trees based on WCGNA in post-surgery 21 day. (O) Correlations of significant modules to cell subsets in Figure 6G and mouse models. (P) Expressions of genes involved in module turquoise and grey in different cell subsets in Figure 6G. (Q) Sankey maps of immune-related GO terms towards modules and cell subsets with corresponding homologous genes to human cells in post-surgery 14 day (R).

In the immune cell population at day 14, we identified MEbrown, MEturquoise, MEred and MEgrey, which were strongly correlated with phenotype and clustering, respectively (Figures 7G-I). MEbrown is mainly in cluster 0, 5, 6, MEturquoise is mainly in cluster 0, 5, 10, 14, MEred is mainly in cluster 3, 9, and MEgrey is distributed in all clusters (Figure J). The corresponding GO-based functions of these modules are : Mainly brown, cell death, MAPK cascade reaction; MEgrey, T cell activation, bone resorption, electron transport chain; MEred, B cell activation; MEturquoise, GO terms related to neutrophils and osteoclasts (Figure K), while the human cell group was corresponding to T cell (h14cluster 0, T cell activation), monocyte (h14cluter 1, 6, cell death), B cell (h14cluster 4, cell death and electron transport chain), B cell (h14cluster 5, B cell activation and differentiation, bone resorption) (Figure L). In non-immune cells, we confirmed that MEpurple, MEturquoise, MEgreen, and MEyellow have strong correlations with phenotype and clustering, respectively (Figures S7F-H). MEpurple is mainly in cluster 0, 8, MEturquoise is in cluster 11, MEgreen is involved in cluster 10, 12 and MEyellow is distributed in cluster 2, 4, 6, 8, 12 (Figures S7I). The functions are that: MEpurple, angiogenesis, Notch signaling pathway, Wnt signaling pathway; MEgreen, vasculature development, extracellular matrix composition; MEturquoise, mitosis; MEyellow, mainly programmed cell death, interleukin I feedback, adipocyte differentiation, EAR1/2 cascade (Figures S7J).

In the immune cell population at 21 days, we confirmed that MEbrown, MEturquoise and MEyellow were strongly correlated with phenotype and clustering, respectively (Figures 7M-O). MEbrown is mainly in cluster 0, 2, 5, 6, MEturquoise is distributed in all clusters and MEyellow is mainly in cluster 9, 15 (Figure P). The corresponding functions of these modules are that: MEbrown, neutrophil immunity; MEturquoise, ribosome biosynthesis, ncRNA transcription, neutrophil migration; MEyellow, B cell activation, complement activation and phagocytosis (Figure Q), while in the corresponding human-based cell population, they were B cell (h21cluster 0, 1, B cell activation and ncRNA processing) and neutrophils (h21cluster 2, neutrophil-mediated immunity) (Figure R). In non-immune cells, we confirmed that MEred, MEturquoise, MEgreen and MEgrey have strong correlations with phenotype and clustering, respectively (Figures S7K-M). MEgreen is mainly in cluster 12, 14, and 17, and MEgrey is in all clusters, MEred is in cluster 0, 1, and MEturquoise is in cluster 14, 15, 17 (Figures S7N). The functions are that: MEgreen, mitosis, DNA replication; MEgrey, osteogenesis, angiogenesis, nerve formation, mineralization, wnt signaling pathway, glucose metabolism, oxidative phosphorylation, electron transport chain; MEred, angiogenesis, chondrocyte differentiation, extracellular matrix; MEturquoise, mast cell activation, phagocytosis, neutrophil migration (Figures S7O).

## Discussion

At present, it is considered that bone repair is a complex process with interaction of various cells in a spatiotemporal manner, such as immune cell, stem cells, endothelial cells, etc. Here, we systematically revealed the cell dynamics and their corresponding functions at different stage of bone repair by using single-cell sequencing, and illustrated the diversity and complementarity of myeloid and lymphoid cells to the bone homeostasis.

Human lymphoid cells were detected to work at day 14 (mainly T cells and NK cells) and 21 (mainly B cells), but not day 7 in the bone repair process. Although they might only bind, but not signal to mouse cytokines, they can respond to the repair signals (*21*) and inflammatory signals. Thus, we can have a sight into the repair reaction from adaptive immune system without signaling to the regenerative tissue. By comparing the single-cell transcription profiles among each sample, we did find that transcription profiles of n7 and cd7 were similar to sc14, but not sc7, which is consistent with the results that bone repair would be accelerated at the early stage in the absence of the adaptive immune system (*8*). On the other side, n21 and cd21 had transcription profiles more similar with sc14 than sc21, suggesting the immune disorder would impact the bone remodeling process (*9*).

Cd8^+^Il-17^+^ T cells and Tregs expand from day 7 to day 14 at the repair site (Figure 2A) and become major sources of Il-17 (Figure 4C) and Tnf (Figure 5A, B) during the bone repair process, respectively. The former may contribute to T cell inflammation and necrotic tissue damage (Figures 4D and 4E). Although the latter may have adopted an effector-like state, they are still more enriched for Treg cell markers (e.g., Foxp3, Ctla4, and Il10) than Tnf–Treg cells. Future work will determine whether Tnf^+^ Treg cells contribute to bone reformation or anti-Tnf resistance (*22*), as well as the role of Cd8^+^ T cell plasticity in bone repair-related inflammation.

Neutrophil signaling was implicated in cytokine resistance via unknown myeloid and stromal cell types (*23*). Here we show that inflammatory monocytes and inflammation-associated fibroblast (IAF)s may mediate resistance via expression of Elane and Ptne, respectively (Figure 5D). In particular, IAFs were enriched in n7 and cd7 groups (Figure 5F). In addition, we identified that basophils phenocopies B cells, which may explain bone remodel complementation. Future work will determine whether IAFs are a robust biomarker of repair response or whether combining neutrophils with inhibition of IAF cytokines and/or receptors can reduce cytokine resistance in immune disorder populations.

IAFs uniquely express Il11, a potential therapeutic target for fibrosis (*24*), suggesting involvement in gut fibrosis. Because they express osteogenesis-associated (Wnt2b^+^) and angiogenesis-associated (Wnt5b^+^) markers (Figures 1I and S3E), IAFs may reflect a distinct fibroblast state. IAFs express several angiogenesis markers, and IAF markers are enriched in angiogenesis of tumors (*25*), suggesting a shared origin and/or state. IAF expansion during bone repair-associated inflammation may affect the bone marrow/bone defect microenvironment. Last, both IAFs and inflammatory monocytes form hubs in the cell-cell interaction network and may affect the proportions of other cells (Figure 6E; Table S5).

Our work provides a framework for using scRNA-seq to understand bone repair process and its immune responses. We identify changes in cell proportions and gene expression with immune disorder state and integrate these to understand mechanisms of cell-cell signaling and drug susceptibility. Finally, we nominate hub genes across loci, predicting their cells of action and putative functions, and assemble them into the core pathways that underlie bone repair.

## Materials and Methods

### REAGENT or RESOURCE

Alizarin red S, Sigma, USA

Calcein, Sigma, USA

Type I Collagenase, Sigma-Aldrich, Cat#C6885

Type II Collagenase

DNA lyase

Tesca Buffer, Solarbio, G0150

### Software and Algorithms

Cellranger R 4.0.5

Seurat 4.0.3

SingleR 1.4.1

Monocle 2.18.0

Igraph 1.2.6

GOstats 2.56.0

ggplot2 3.3.5

ggalluvial 0.12.3

genefilter 1.72.1

dplyr 1.0.7

destiny 3.4.0

celldex 1.0.0

GO.db 3.12.1

Limma 3.46.0

org.Hs.eg.db 3.12.0

org.Mm.eg.db 3.12.0

WGCNA 1.70-3

Contact for reagent and resource sharing

Further information and requests for resources and reagents should be directed to and will be fulfilled by the lead contact Xinquan Jiang

### Experimental model and subject details

#### Mice

ICR mice were purchased from Phenotek Biotechnology (Shanghai, China).

All procedures were conducted under animal care protocols approved by the Animal Research Committee of the Ninth People’s Hospital affiliated to Shanghai Jiao Tong University, School of Medicine[SH9H-2019-A167-1].

#### Generation of NPSG and CD34^+^NPSG mice

NPSG:NPSG mice (NOD-*Prkdc*^scid^ *IL2rg*^null^/Pnk) were modified by CRISPR/Cas9 gene editing system. Based on NOD strain mice, *Prkdc* gene knockout firstly and then *IL2rg* gene was knocked out. Genotype and phenotype of NPSG mice were verified by PCR, Flow Cytometry and *in vivo* experiment. Highly defection of T, B and NK cells is suitable for tumor research and humanized immune system reconstruction.

CD34^+^ NPSG:NPSG mice were developed at Phenotek Biotechnology (Shanghai, China) by backcrossing a complete null mutation at the *SCID* and *Il2rg* locus onto the NOD strain. Purified human CD34^+^ hematopoietic stem cells (HPSCs) were purchased from Milestone Biotechnologies (Shanghai, China). CD34^+^ humanized NPSG mice were generated according to this guideline. In brief, CD34^+^ humanized NPSG mice were generated by intravenous injection of 2.5 × 10^4^ human CD34^+^ HPSCs into 3 weeks-old female NPSG mice, 4h post 1.4Gy total body irradiation using the RS2000 Pro irradiator (Rad Source, Buford, GA, USA). The engraftment levels of hCD45^+^ cells were determined 12 weeks post-HPSCs transplantation by flow cytometric quantification of peripheral blood human CD45^+^ cells. CD34^+^ humanized NPSG mice that had over 25% human CD45^+^ cells in the peripheral blood were considered as engrafted and humanized.

#### Mice bone hole drilling model

A bone hole drilling model was used to investigate the physiological bone repair. The mice were anaesthetized under inhalation anesthesia with isoflurane. A 3-mm diameter critical-sized defects were created using a trephine bur (*26*). Following surgery, the soft tissues were repositioned and sutured. The mice were injected with calcein at 7day, and Alizarin red S at 14day post-surgery. The mice were injected with sacrificed at 7day, 14day and 21day post-surgery, and the soft and hard tissues in the bone repair region were harvested for scRNA-sequencing and the whole skull were harvested for micro-CT assay.

#### Isolation of repaired tissue from drilled bone hole

The tissues in the bone repair region were minced separately with razor blades, resuspended in 3000 U/mL type II collagenase (Sigma-Aldrich, Cat#C6885) digestion buffer supplemented with 100 U/mL DNase I (Worthington, Cat#NC9199796) and incubated at 37 C for 40 min under constant agitation. The supernatant was filtered through a 70 mm nylon mesh and quenched with staining media (2% fetal bovine serum (FBS), in phosphate-buffered saline (PBS), GIBCO, Cat#C14190500BT). Digestion was repeated twice more prior to centrifugation at 200 g at 4 C followed by resuspension in staining media (*12*).

#### Micro-CT assay

The skull was harvested and fixed in 4% paraformaldehyde for 24h. Then the samples were imaged by micro-CT (SkyScan 1176, Bruker, USA) to analyze the newly formed bone. The scanning parameters were set according to our previous study (*26*). Briefly, the samples were scanned in high-resolution scanning mode (pixel matrix, 1024 × 1024; voxel size, 20 μm; slice thickness, 20 μm). After scanning, three-dimensional images were reconstructed with GEHC MicroView software. The bone mineral density (BMD) and the percentage of new bone volume relative to tissue volume (BV/TV) were measured using auxiliary software (Scanco Medical AG, Switzerland). The volumes of the regenerated bone were determined using DataViewer software (Sky-Scan) and the CTAn program (SkyScan).

#### Sequential fluorescent labeling

A polychrome sequential fluorescent labeling method was used to visualize the process of new bone formation and mineralization. At 7, 14, and 21day after surgery, 25 mg/kg tetracycline hydrochloride (Sigma, USA), 30 mg/kg alizarin red S (Sigma, USA), and 20 mg/kg calcein (Sigma, USA) were intraperitoneally administered.

#### Surface morphology and histological analysis

For subcutaneous implantation, the samples together with the surrounding subcutaneous tissue were harvested with scissors and scalpels. The collected samples were fixed in 4% paraformaldehyde overnight. All the implants were then carefully stripped from the surrounding tissues. The implants were scanned with SEM, and the corresponding surrounding tissues were embedded in paraffin wax. For each sample, 6 μm sections were cut off and mounted onto glass slide for hematoxylin & eosin (H&E) staining. Images were obtained and analyzed with a Motic easy scan digital slide scanner (MOTIC, China).

#### Single Cell RNA-seq

Single cells were encapsulated into emulsion droplets using Chromium Controller (10x Genomics). scRNA-seq libraries were constructed using Chromium Single Cell 30 v2 Reagent Kit according to the manufacturer’s protocol. Briefly, post sorting sample volume was decreased and cells were examined under a microscope and counted with a hemocytometer. Cells were then loaded in each channel with a target output of $4,000 cells. Reverse transcription and library preparation were performed on C1000 Touch Thermal cycler with 96-Deep Well Reaction Module (Bio-Rad). Amplified cDNA and final libraries were evaluated on an Agilent BioAnalyzer using a High Sensitivity DNA Kit (Agilent Technologies). Individual libraries were diluted to 4nM and pooled for sequencing. Pools were sequenced with 75 cycle run kits (26bp Read1, 8bp Index1 and 55bp Read2) on the NextSeq 500 Sequencing System (Illumina) to $70%–80% saturation level.

#### Single-cell sequencing data processing

ScRNA-Seq data were demultiplexed, aligned to the mouse genome, version mm10, and UMI-collapsed with the Cellranger toolkit (version 2.0.1, 10X Genomics). We excluded cells with fewer than 500 detected genes (where each gene had to have at least one UMI aligned) for mouse and 1000 detected genes for human. Gene expression was represented as the fraction of its UMI count with respect to total UMI in the cell and then multiplied by 10,000. We denoted it by TP10K – transcripts per 10K transcripts.

#### Dimensionality reduction

We performed dimensionality reduction using gene expression data for a subset of variable genes. The variable genes were selected based on dispersion of binned variance to mean expression ratios using *FindVariableGenes* function of Seurat package (*27*) followed by filtering of mitochondrial genes. Next, we performed principal component analysis (PCA) and reduced the data to the top 50 PCA components (number of components was chosen based on standard deviations of the principal components – in a plateau region of an ‘‘elbow plot’’).

#### Clustering and sub-clustering

We used graph-based clustering of the PCA reduced data with the Louvain Method after computing a shared nearest neighbor graph (*28*). We visualized the clusters on a 2D map produced with t-distributed stochastic neighbor embedding (t-SNE). For sub-clustering, we applied the same procedure of finding variable genes, dimensionality reduction, and clustering to the restricted set of data (usually restricted to one initial cluster).

#### GO-based Gene Enrichment Analysis

The functional enrichment analysis was performed following the previous study (*19*). The expression of individual genes was defined as altered when comparison of the average normalized signal intensities using the Bioconductor package Genefilter gave a value of p<0.05 in Welch’s ANOVA. To investigate the ontology of marker genes, enriched expression of GO terms was assessed and confirmed with the Hypergeometric Test15) and Gene Set Enrichment Analysis) using the Bioconductor pack-age GOstats/GSEABase. These programs determine which GO terms identified among the lists of affected genes are statistically over- or underrepresented, as compared with the GO terms represented in the microarray as a whole. Hierarchical clustering was performed using Ward’s method to calculate the linkage distances based on the correlation coefficient between clusters or GO terms.

#### Statistical Analysis

Experiments were conducted with at least three samples for three independent times. All the data were expressed as the mean±standard deviation (SD). The data were analyzed using R 4.0.5 and the statistically significant differences (p) between groups were identified using one-way analysis of the variance and Tukey’s multiple comparison tests. Probability values of <0.05 were considered significant. The differences were expressed at * p<0.05, ** p<0.01 and *** p<0.001.

## Acknowledgments

Thanks to Mr. Minghui Teng and Mrs. Huiying Tao from Shanghai Personalbio Technology Co. Ltd. for the cooperation of single-cell RNA sequencing and data uploading. Thanks to Mr. Shaohua Deng from Phenotek Biotechnology (Shanghai, China) for NPSG mice supporting.

## Funding

1. the National Natural Science Foundation of China (81921002, Jiang. X; 32000812, Shi. J;81670968, Chang. Q)
2. Innovative research team of high-level local universities in Shanghai, SHSMU-ZLCX20212400 (Jiang. X)
3. The National Key Research and Development Program of China, 2016YFC1102900 (Jiang. X)

## Author contributions

Jiang.X, Chang.Q and Shi.J. conceived the concept of the study and provided funding support. Shi.J. and Zhang.M. generated the cranial defect model. Shi.J., Wang.J. and Jiang.F. performed histological analysis and conducted the part of sample preparation and characterization. Shi.J., Yin.S, Lin.S. and Wu.X. provided the micro-CT technology support, image processing and essential data analysis. Shen.L., Yang.J, Wen. J., Gu. X, Yang. R. and Zhang.W. provided essential technology support and experimental materials. Shi.J. performed single cell-sequencing data analysis. Shi.J. and Wang.J. drafted the manuscript. Jiang. X and Chang. Q reviewed the manuscript; Shi. J. and Wang.J. integrated the suggestions of all authors and finished the manuscript.

## Competing interests

The authors declare no competing interests.

## Data and materials availability

The scRNA-seq data generated in this study are deposited in GEO (GSE178949, https://www.ncbi.nlm.nih.gov/geo/query/acc.cgi? acc=GSE178949).

## Figures and Tables

**Figure S1.**
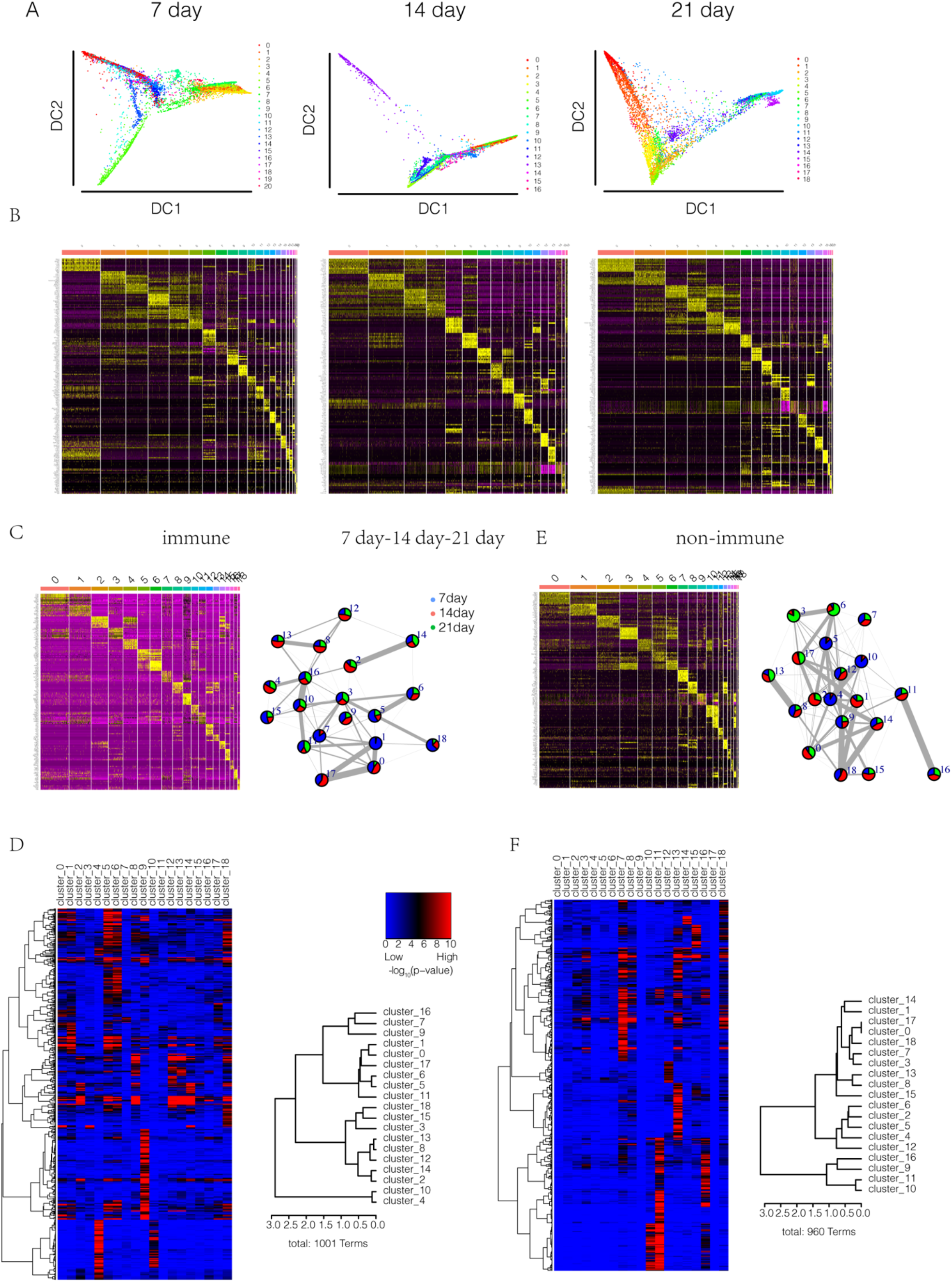
(A) Diffusion maps of cells colored by cell subsets in different time points. (B) Marker genes in different cell subsets. (C) Marker genes in different immune cell subsets and relatedness (edges, width indicates strength) between clusters (A) (nodes, colored as mouse models) based on cluster graph abstraction. (D) GO heatmap and clustering dendrogram of cell subsets in C with GO terms along the horizontal axis. The heatmap was generated across all cell subsets. P values were calculated by the hypergeometric test and denoted by -log_10_P; overrepresented GO terms were shown in each cell subsets and colored by red and blue in Extend Figure-6Q; distance method: correlation; agglomeration method: Ward’s method. Cluster dendrogram of cell subsets in C based on the patterns of GO enrichment; distance method: correlation; agglomeration method: Ward’s method. The linkage distance is the measure of similarity between cell subsets, with increasing distance representing decreasing degrees of similarity between samples. (E) Marker genes in different non-immune cell subsets and relatedness (edges, width indicates strength) between clusters (nodes, colored as mouse models) based on cluster graph abstraction. (F) GO heatmap and clustering dendrogram of cell subsets in E with GO terms along the horizontal axis. The heatmap was generated across all cell subsets. P values were calculated by the hypergeometric test and denoted by -log_10_P; overrepresented GO terms were shown in each cell subsets and colored by red and blue in Extend Figure-6Q; distance method: correlation; agglomeration method: Ward’s method. Cluster dendrogram of cell subsets in E based on the patterns of GO enrichment; distance method: correlation; agglomeration method: Ward’s method. The linkage distance is the measure of similarity between cell subsets, with increasing distance representing decreasing degrees of similarity between samples.

**Figure S2.**
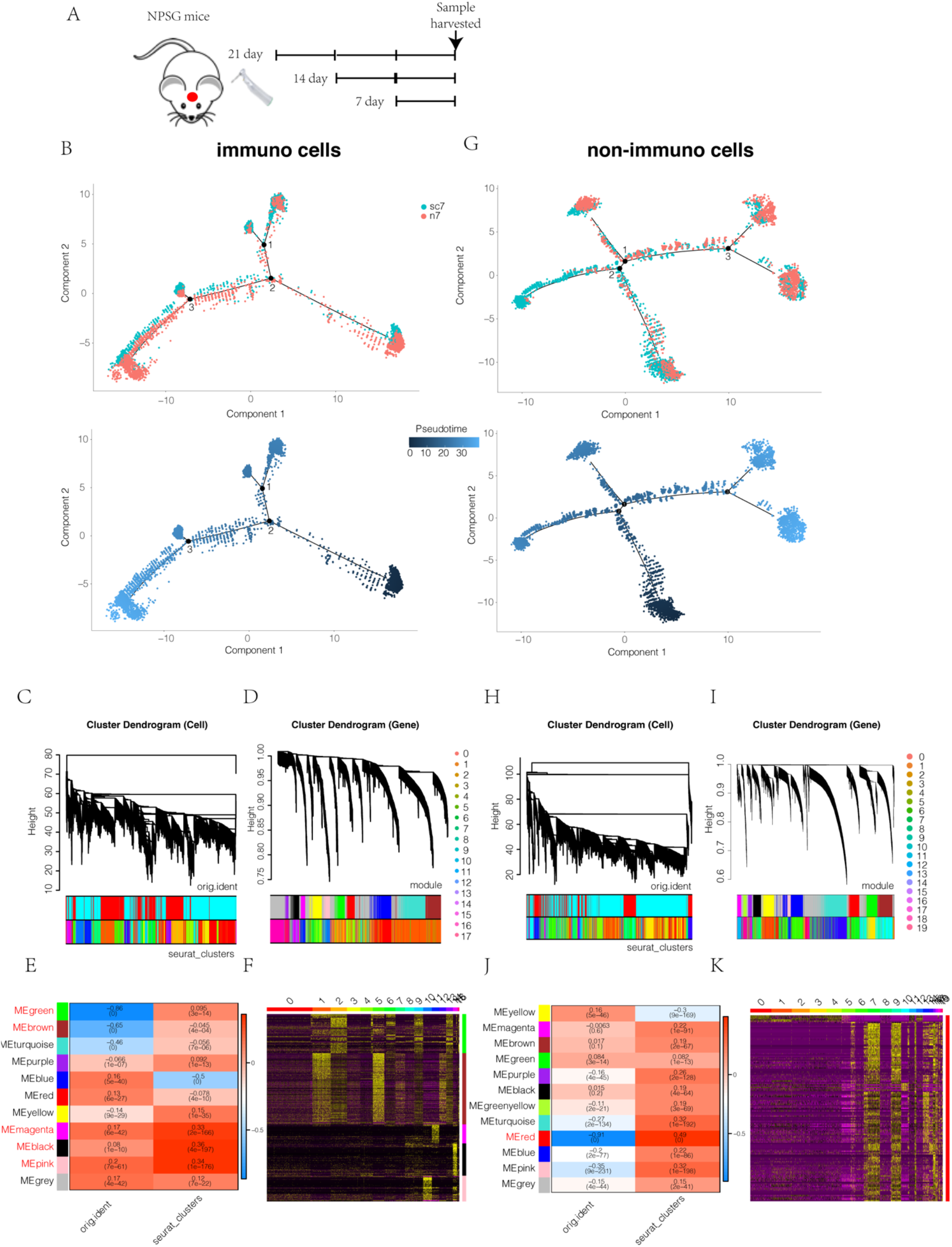
(A) Study design. (B) Pseudotime analysis of immune cells colored by pseudotime and mouse models. (C) Cell and (D) gene trees based on WCGNA in post-surgery 7 day. (E) Correlations of significant modules to cell subsets in Figure 2A and mouse models. (E) (F) Expressions of genes involved in modules highlighted in C in different cell subsets in Figure 2A. (G) Pseudotime analysis of immune cells colored by pseudotime and mouse models. (H) Cell and (I) gene trees based on WCGNA in post-surgery 7 day. (J) Correlations of significant modules to cell subsets in Figure 2G and mouse models. (K) Expressions of genes involved in modules highlighted in C in different cell subsets in Figure 2G.

**Figure S3.**
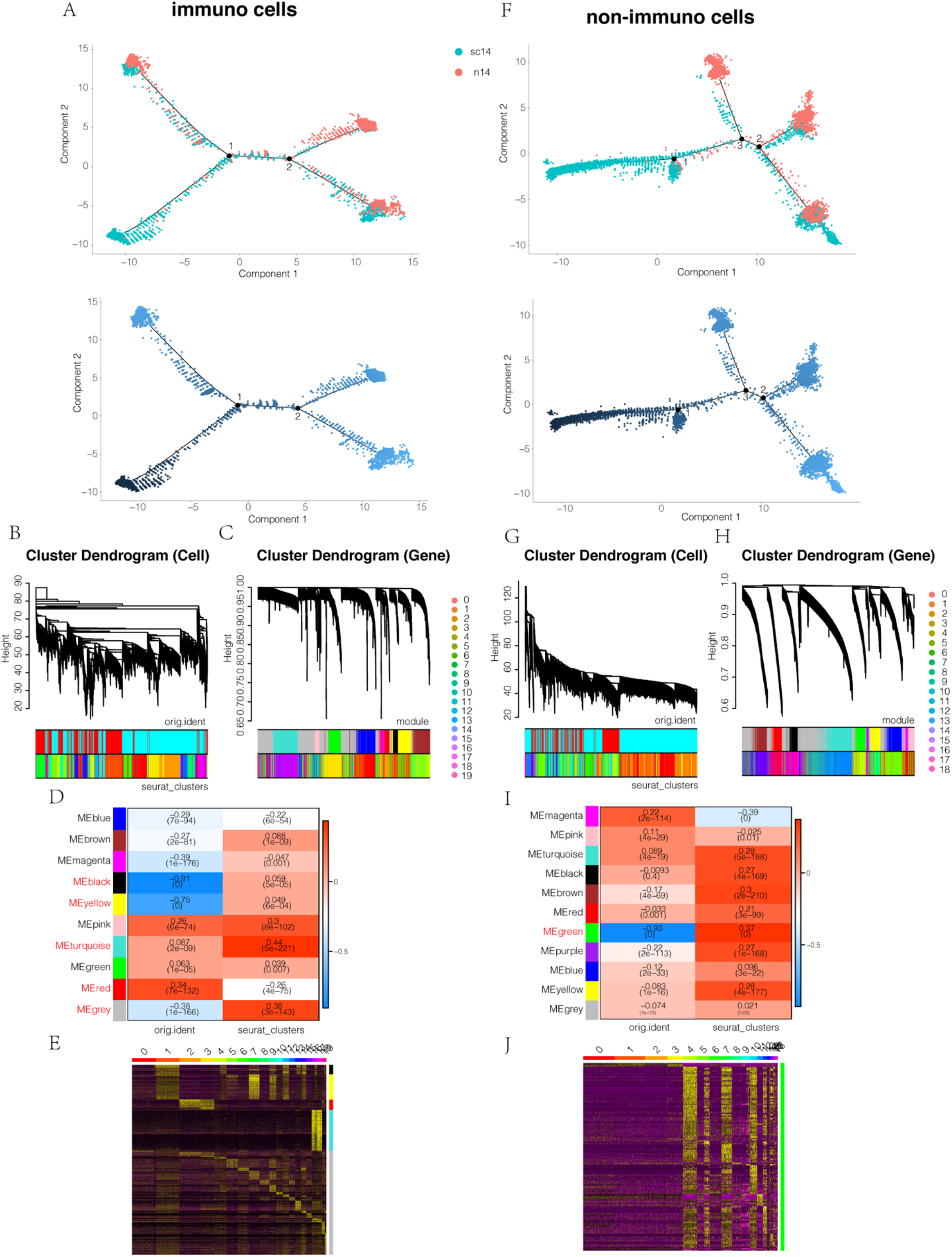
(A) Pseudotime analysis of immune cells colored by pseudotime and mouse models. (B) Cell and (C) gene trees based on WCGNA in post-surgery 14 day. (D) Correlations of significant modules to cell subsets in Figure 3A and mouse models. (E) Expressions of genes involved in modules highlighted in C in different cell subsets in Figure 2A. (F) Pseudotime analysis of immune cells colored by pseudotime and mouse models. (G) Cell and (H) gene trees based on WCGNA in post-surgery 14 day. (I) Correlations of significant modules to cell subsets in Figure 3G and mouse models. (J) Expressions of genes involved in modules highlighted in C in different cell subsets in Figure 3G.

**Figure S4.**
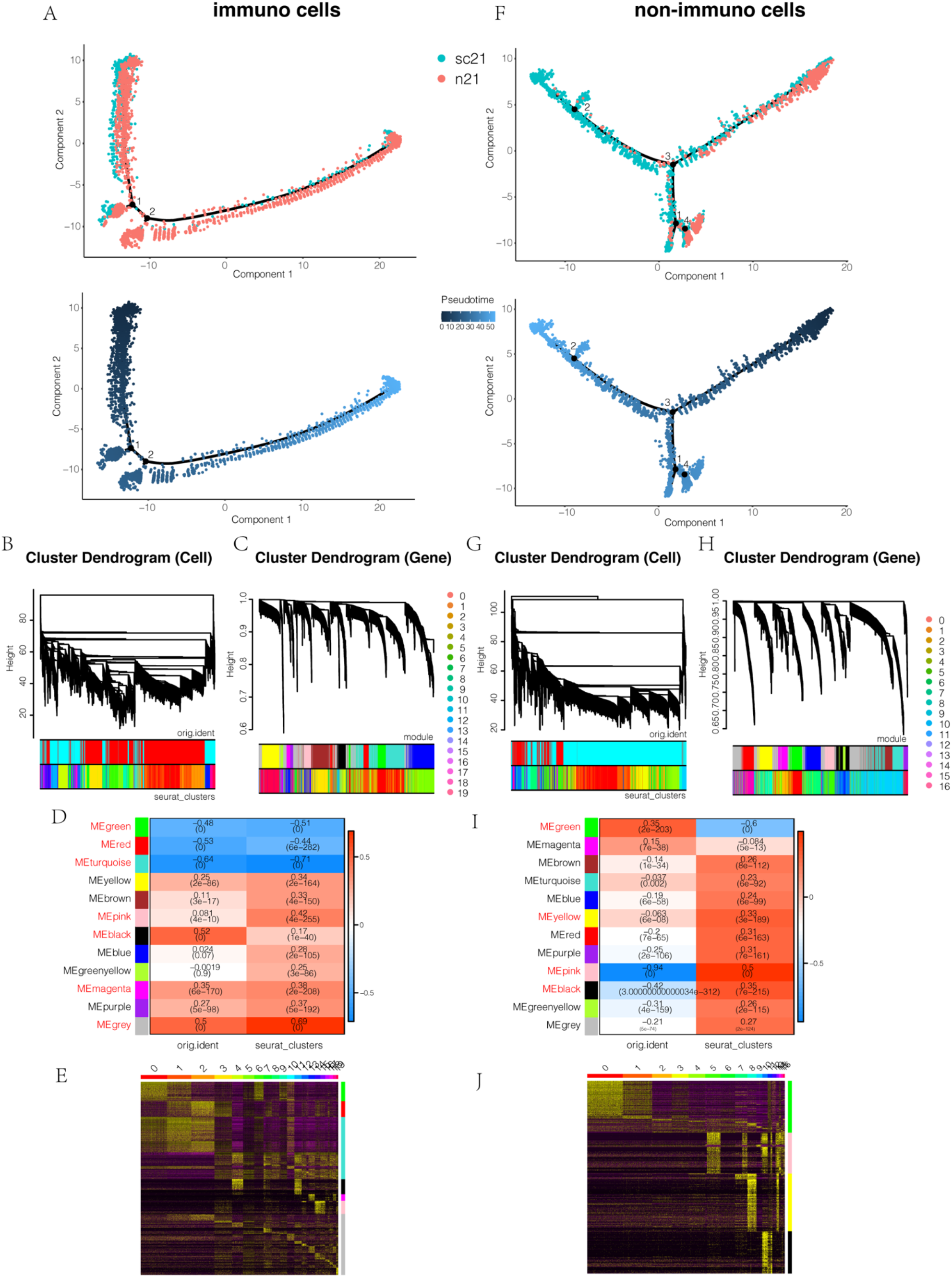
(B) Pseudotime analysis of immune cells colored by pseudotime and mouse models. (B) Cell and (C) gene trees based on WCGNA in post-surgery 21 day. (D) Correlations of significant modules to cell subsets in Figure 4A and mouse models. (D) (E) Expressions of genes involved in modules highlighted in C in different cell subsets in Figure 2A. (F) Pseudotime analysis of immune cells colored by pseudotime and mouse models. (G) Cell and (H) gene trees based on WCGNA in post-surgery 21 day. (I) Correlations of significant modules to cell subsets in Figure 4G and mouse models. (J) Expressions of genes involved in modules highlighted in C in different cell subsets in Figure 4G.

**Figure S5.**
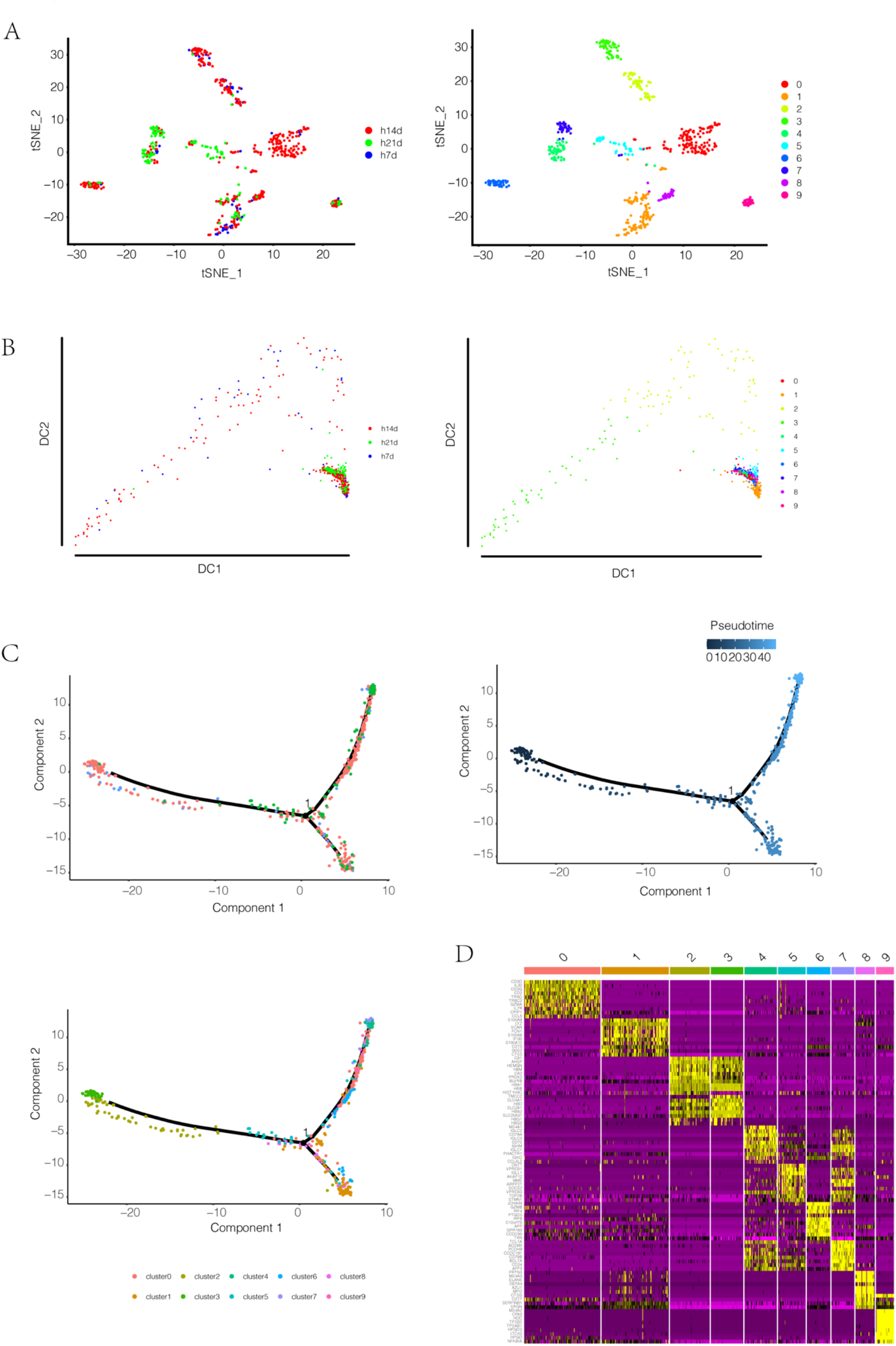
(A) Cell census at bone reformation site colored by human cell subsets and post-surgery time points from t-SNE, diffusion map analysis (B) and pseudotim analysis (C) colored by post-surgery time points, pseudotime and cell subsets. (C) Marker genes in cell subsets of A.

**Figure S6.**
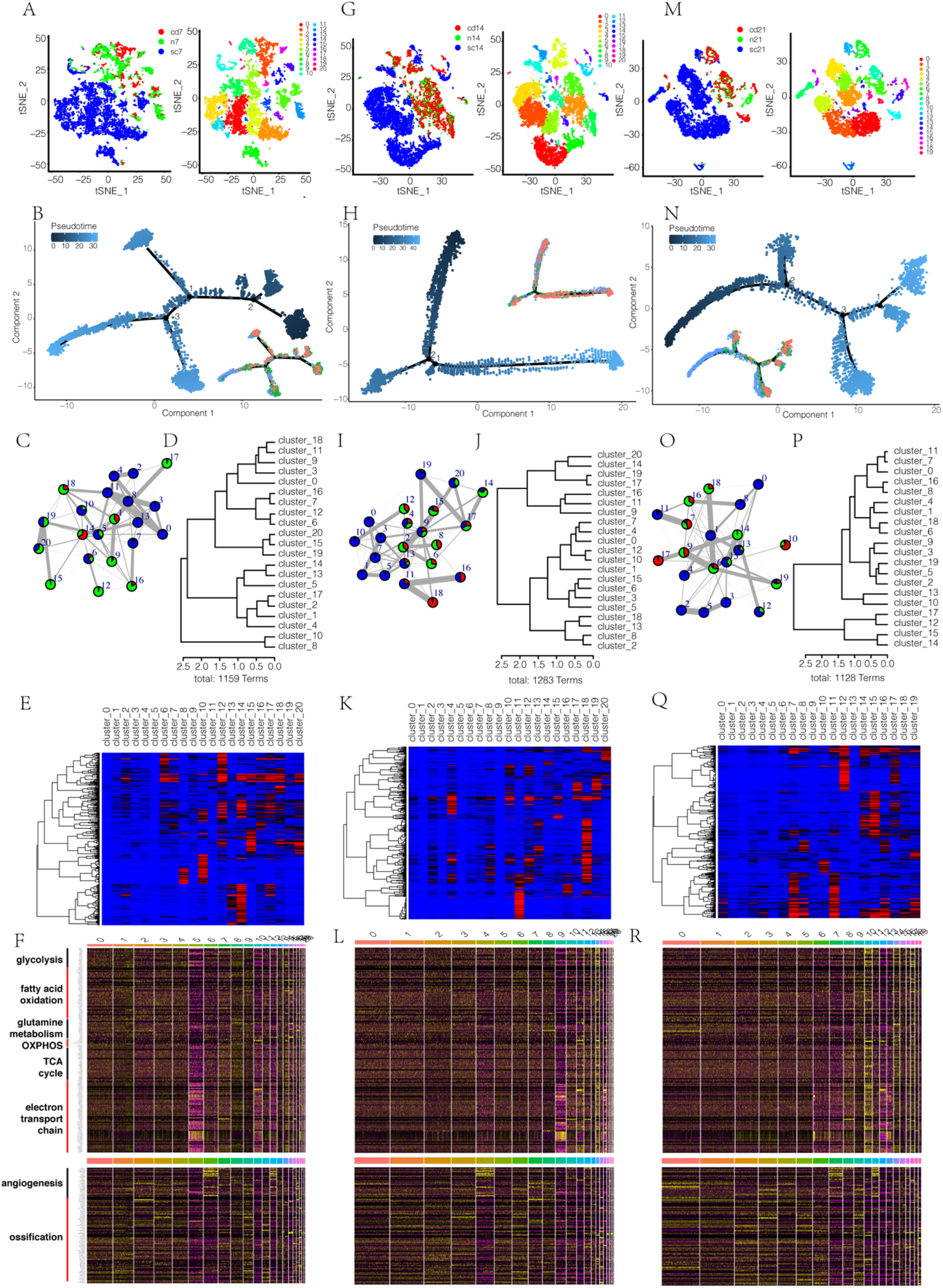
(A) Cell census at bone reformation site colored by cell subsets and different mouse models at post-surgery 7 day from t-SNE and diffusion map analysis. (A) (B) Pseudotime analysis of cells in A colored by pseudotime and mouse models. (C) Relatedness (edges, width indicates strength) between clusters (nodes, colored as mouse models) based on cluster graph abstraction. (D) Cluster dendrogram of cell subsets in C based on the patterns of GO enrichment; distance method: correlation; agglomeration method: Ward’s method. The linkage distance is the measure of similarity between cell subsets, with increasing distance representing decreasing degrees of similarity between samples. (E) GO heatmap and clustering dendrogram of cell subsets in A with GO terms along the horizontal axis. The heatmap was generated across all cell subsets. P values were calculated by the hypergeometric test and denoted by -log_10_P; overrepresented GO terms were shown in each cell subsets and colored by red and blue in Extend Figure-6E; distance method: correlation; agglomeration method: Ward’s method. (F) Marker genes related to metabolisms in different cell subsets in A. (G) Cell census at bone reformation site colored by cell subsets and different mouse models at post-surgery 14 day from t-SNE and diffusion map analysis. (H) Pseudotime analysis of cells in G colored by pseudotime and mouse models. (I) Relatedness (edges, width indicates strength) between clusters (nodes, colored as mouse models) based on cluster graph abstraction. (J) Cluster dendrogram of cell subsets in I based on the patterns of GO enrichment; distance method: correlation; agglomeration method: Ward’s method. The linkage distance is the measure of similarity between cell subsets, with increasing distance representing decreasing degrees of similarity between samples. (K) GO heatmap and clustering dendrogram of cell subsets in G with GO terms along the horizontal axis. The heatmap was generated across all cell subsets. P values were calculated by the hypergeometric test and denoted by -log_10_P; overrepresented GO terms were shown in each cell subsets and colored by red and blue in Extend Figure-6K; distance method: correlation; agglomeration method: Ward’s method. (L) Marker genes related to metabolisms in different cell subsets in G. (M) Cell census at bone reformation site colored by cell subsets and different mouse models at post-surgery 21 day from t-SNE and diffusion map analysis. (N) Pseudotime analysis of cells in M colored by pseudotime and mouse models. (O) Relatedness (edges, width indicates strength) between clusters (nodes, colored as mouse models) based on cluster graph abstraction. (P) Cluster dendrogram of cell subsets in M based on the patterns of GO enrichment; distance method: correlation; agglomeration method: Ward’s method. The linkage distance is the measure of similarity between cell subsets, with increasing distance representing decreasing degrees of similarity between samples. (Q) GO heatmap and clustering dendrogram of cell subsets in M with GO terms along the horizontal axis. The heatmap was generated across all cell subsets. P values were calculated by the hypergeometric test and denoted by -log_10_P; overrepresented GO terms were shown in each cell subsets and colored by red and blue in Extend Figure-6Q; distance method: correlation; agglomeration method: Ward’s method. (R) Marker genes related to metabolisms in different cell subsets in M.

**Figure S7.**
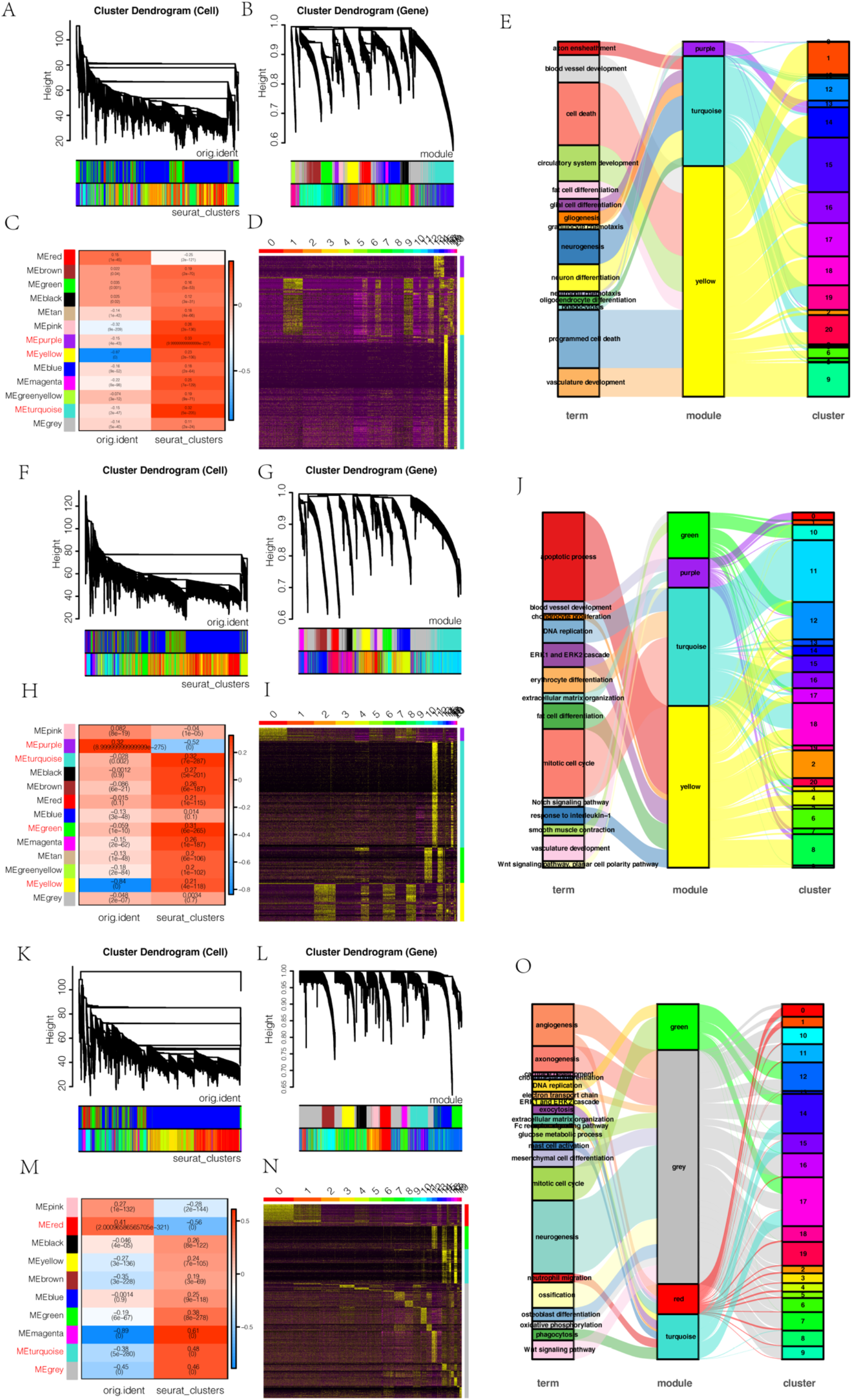
(A) Cell and (B) gene trees based on WCGNA in post-surgery 7 day. (C) Correlations of significant modules to cell subsets in Figure 6A and mouse models. (D) Expressions of genes involved in module turquoise and grey in different cell subsets in Figure 6A. (E) Sankey maps of immune-related GO terms towards modules and cell subsets with corresponding homologous genes to human cells in post-surgery 7 day (F). (G) Cell and (H) gene trees based on WCGNA in post-surgery 14 day. (I) Correlations of significant modules to cell subsets in Figure 6G and mouse models. (J) Expressions of genes involved in module turquoise and grey in different cell subsets in Figure 6G. (K) Sankey maps of immune-related GO terms towards modules and cell subsets with corresponding homologous genes to human cells in post-surgery 14 day (L). (M) Cell and (N) gene trees based on WCGNA in post-surgery 21 day. (O) Correlations of significant modules to cell subsets in Figure 6G and mouse models. (P) Expressions of genes involved in module turquoise and grey in different cell subsets in Figure 6G. (Q) Sankey maps of immune-related GO terms towards modules and cell subsets with corresponding homologous genes to human cells in post-surgery 14 day (R).

## Notes

### Competing Interest Statement

The authors have declared no competing interest.

## References

1. G. Eraslan et al., Single-nucleus cross-tissue molecular reference maps toward understanding disease gene function. Science 376, eabl4290 (2022).

2. C. Dominguez Conde et al., Cross-tissue immune cell analysis reveals tissue-specific features in humans. Science 376, eabl5197 (2022).

3. G. Hermeren, The ethics of regenerative medicine. Biol Futur 72, 113–118 (2021).

4. A. Salhotra, H. N. Shah, B. Levi, M. T. Longaker, Mechanisms of bone development and repair. Nat Rev Mol Cell Biol 21, 696–711 (2020).

5. E. C. Kruijt Spanjer, G. K. P. Bittermann, I. E. M. van Hooijdonk, A. Rosenberg, D. Gawlitta, Taking the endochondral route to craniomaxillofacial bone regeneration: A logical approach? J Craniomaxillofac Surg 45, 1099–1106 (2017).

6. S. Yin, W. Zhang, Z. Zhang, X. Jiang, Recent Advances in Scaffold Design and Material for Vascularized Tissue-Engineered Bone Regeneration. Adv Healthc Mater 8, e1801433 (2019).

7. L. Claes, S. Recknagel, A. Ignatius, Fracture healing under healthy and inflammatory conditions. Nat Rev Rheumatol 8, 133–143 (2012).

8. D. Toben et al., Fracture healing is accelerated in the absence of the adaptive immune system. J Bone Miner Res 26, 113–124 (2011).

9. A. E. Rapp et al., Fracture Healing Is Delayed in Immunodeficient NOD/scidIL2Rgammacnull Mice. PLoS One 11, e0147465 (2016).

10. C. K. F. Chan et al., Identification of the Human Skeletal Stem Cell. Cell 175, 43–56 e21 (2018).

11. C. C. Lee, N. Hirasawa, K. G. Garcia, D. Ramanathan, K. D. Kim, Stem and progenitor cell microenvironment for bone regeneration and repair. Regen Med 14, 693–702 (2019).

12. N. Baryawno et al., A Cellular Taxonomy of the Bone Marrow Stroma in Homeostasis and Leukemia. Cell 177, 1915–1932 e1916 (2019).

13. N. Yang, Y. Liu, The Role of the Immune Microenvironment in Bone Regeneration. Int J Med Sci 18, 3697–3707 (2021).

14. Z. Li, J. A. Helms, Drill Hole Models to Investigate Bone Repair. Methods Mol Biol 2221, 193–204 (2021).

15. Z. S. Ai-Aql, A. S. Alagl, D. T. Graves, L. C. Gerstenfeld, T. A. Einhorn, Molecular mechanisms controlling bone formation during fracture healing and distraction osteogenesis. J Dent Res 87, 107–118 (2008).

16. G. Schiebinger et al., Optimal-Transport Analysis of Single-Cell Gene Expression Identifies Developmental Trajectories in Reprogramming. Cell 176, 928–943 e922 (2019).

17. C. Trapnell et al., The dynamics and regulators of cell fate decisions are revealed by pseudotemporal ordering of single cells. Nat Biotechnol 32, 381–386 (2014).

18. F. A. Wolf et al., PAGA: graph abstraction reconciles clustering with trajectory inference through a topology preserving map of single cells. Genome Biol 20, 59 (2019).

19. N. Liu et al., Effects of a Tricaprylin Emulsion on Anti-glomerular Basement Membrane Glomerulonephritis in Rats: In Vivo and in Silico Studies. Biol Pharm Bull 38, 1175–1184 (2015).

20. Y. Iimori, M. Morioka, S. Koyamatsu, N. Tsumaki, Implantation of Human-Induced Pluripotent Stem Cell-Derived Cartilage in Bone Defects of Mice. Tissue Eng Part A 27, 1355–1367 (2021).

21. N. C. Walsh et al., Humanized Mouse Models of Clinical Disease. Annu Rev Pathol 12, 187–215 (2017).

22. R. Atreya et al., Antibodies against tumor necrosis factor (TNF) induce T-cell apoptosis in patients with inflammatory bowel diseases via TNF receptor 2 and intestinal CD14(+) macrophages. Gastroenterology 141, 2026–2038 (2011).

23. N. R. West et al., Oncostatin M drives intestinal inflammation and predicts response to tumor necrosis factor-neutralizing therapy in patients with inflammatory bowel disease. Nat Med 23, 579–589 (2017).

24. S. Schafer et al., IL-11 is a crucial determinant of cardiovascular fibrosis. Nature 552, 110–115 (2017).

25. K. Micka-Michalak, T. Biedermann, E. Reichmann, M. Meuli, A. S. Klar, Induction of angiogenic and inflammation-associated dermal biomarkers following acute UVB exposure on bio-engineered pigmented dermo-epidermal skin substitutes in vivo. Pediatr Surg Int 35, 129–136 (2019).

26. M. Zhang et al., Recapitulation of cartilage/bone formation using iPSCs via biomimetic 3D rotary culture approach for developmental engineering. Biomaterials 260, 120334 (2020).

27. Y. Hao et al., Integrated analysis of multimodal single-cell data. Cell 184, 3573–3587 e3529 (2021).

28. R. Satija, J. A. Farrell, D. Gennert, A. F. Schier, A. Regev, Spatial reconstruction of single-cell gene expression data. Nat Biotechnol 33, 495–502 (2015).

